# The *Drosophila* maternal-effect gene *abnormal oocyte* (*ao*) does not repress histone gene expression

**DOI:** 10.1101/2024.09.17.613536

**Authors:** Risa Takenaka, Eric H. Albanese, Sierra M. Simmerman, Shilpi Verghese, Mandalay A.E. Maddox, Aida Flor de la Cruz, Janet M. Young, Casey A. Schmidt, Leila E. Rieder, Harmit S. Malik

**Author notes:** Correspondence should be addressed to: C. A. S., 746 High St., Easton, PA 18042 USA, L. E. R., 1510 Clifton Rd. NE, Atlanta, GA 30012 USA, H. S. M., 1100 Fairview Avenue N. A2-205, Seattle, WA 98109 USA.

## Abstract

The *abnormal oocyte* (*ao*) gene of *Drosophila melanogaster* is a maternal-effect lethal gene previously identified as encoding a transcriptional regulator of core histones. However, background genetic mutations in existing *ao* mutant strains could compromise their utility in manipulating histone levels. To distinguish the true *ao* phenotype from background effects, we created two new *ao* reagents: a CRISPR/Cas9-mediated knockout of the *ao* allele for genetic and molecular analyses and an epitope-tagged *ao* allele for cytological experiments. Using these reagents, we confirm previous findings that loss of *ao* causes maternal-effect lethality, which can be rescued by either a decrease in the histone gene copy number or by Y chromosome heterochromatin. Our data indicate that *ao* genetically interacts with the heterochromatin, as previously suggested. However, contrary to a prior study, we find neither Ao localization to histone genes nor *ao* repression of core histone transcript levels. Thus, the molecular basis for *ao*-associated maternal-effect lethality remains unknown.

**Article Summary:** A series of foundational papers established that *abnormal oocyte* (*ao*), a euchromatic maternal-effect lethal gene, interacts with heterochromatin and the histone multigene cluster to dictate embryonic viability in *D. melanogaster.* An earlier report argued that *ao* encodes a protein that localizes to and represses histone gene expression, thereby connecting histone gene overexpression with *ao* mutant maternal-effect lethality. Using new reagents for genetics and cytology, we recapitulate findings that *ao* encodes a maternal-effect lethal gene, whose loss is ameliorated by excess heterochromatin or loss of histone genes. However, we find that *ao* does not affect histone gene expression. Thus, how *ao* loss causes maternal-effect lethality remains unknown.

## Introduction

In 1965, Larry Sandler and colleagues collected flies from *Drosophila melanogaster* populations near Rome, Italy, to screen for recessive mutations affecting meiosis. One of the isolated mutants produced excess female offspring when mated to males carrying an attached X^Y chromosome (Sandler *et al*. 1968; Sandler 1970). Sandler named this mutant *abnormal oocyte* (*abo*, recently renamed *ao*) for its aberrant sex-ratio phenotype (Sandler 1970). Subsequent analyses by Sandler showed that *ao* was one of five maternal-effect, embryonic semi-lethal genes located on the left arm of the 2^nd^ chromosome (Sandler 1977). These five genes shared the unusual property that offspring survival from homozygous-mutant mothers was directly affected by the amount of heterochromatin on X and Y chromosomes in the zygote (Sandler 1977). These genes promised to reveal the mechanistic basis of genetic interactions between euchromatin, heterochromatin, and embryonic viability.

Research on *ao* in the following two decades bolstered Sandler’s initial observation that the viability of offspring from *ao* mutant mothers could be rescued by increasing the dosage of certain heterochromatic regions on the X, Y, and 2^nd^ chromosomes (Parry and Sandler 1974; Sandler 1977; Haemer 1978; Yedvobnick *et al*. 1980; Pimpinelli *et al*. 1985; Tomkiel *et al*. 1991). These regions, located on the distal heterochromatin on the X, the long and short arms of the Y (the *Drosophila* Y chromosome is entirely heterochromatic), and the centromeric heterochromatin on the right arm of the 2^nd^ chromosome, were named AO heterochromatic elements (Pimpinelli *et al*. 1985).

The mechanistic relationship between the *ao* mutation and AO heterochromatin remained unclear. Sandler initially hypothesized that an increase in the ribosomal DNA (rDNA) copy number (*i.e.,* the number of rDNA repeats at the locus) was responsible for AO heterochromatin’s amelioration of the maternal-effect lethality. He based this hypothesis on the X- and Y-chromosomal location of the rDNA locus in *D. melanogaster* and the fact that *ao*-associated maternal-effect lethality was lower at 19.5°C (where flies develop more slowly) than at 25.5°C. Indeed, a subsequent study found that *ao* flies maintained as homozygous mutants developed an expansion of the rDNA locus, which alleviated the maternal-effect lethality (Krider and Levine 1975). Later studies observed the same suppressor phenotype in *ao* flies kept in homozygote stocks. However, while some found additional evidence implicating rDNA (Krider *et al*. 1979; Graziani *et al*. 1981), others disputed that rDNA copy number was the cause of the suppressor phenotype (Yedvobnick *et al*. 1980; Pimpinelli *et al*. 1985; Sullivan and Pimpinelli 1986; Cavaliere *et al*. 1991). Furthermore, it remained unclear whether AO heterochromatin rescues the *ao* mutation directly (*i.e*., both *ao* and AO produce the same product) or indirectly *(i.e.,* AO produces a different product than *ao* but performs a rescue function).

Despite over two decades of research, it wasn’t until 1995 that the *ao* mutation was mapped to the cytogenetic locus 32C on the 2^nd^ chromosome of *D. melanogaster* (Tomkiel *et al*. 1995). The genetic unmasking of *ao* took advantage of two *ao* mutants (*ao^1^*, the strain isolated from the Roman fruit-market flies, and *ao^2^*, a *P-*element-induced allele) and a transgenic rescue construct (Tomkiel *et al*. 1995). In 2001, the identity of *ao* was revealed as the gene *CG6093* (Berloco *et al*. 2001), which is the *D. melanogaster* ortholog of the *de-etiolated* or *DET1* gene, first characterized in *Arabidopsis thaliana* but later shown to be present as single-copy orthologs in most plant and animal genomes (Chory *et al*. 1989; Berloco *et al*. 2001).

This study also proposed a molecular mechanism underlying *ao*’s maternal-effect lethality (Berloco *et al*. 2001). It showed that *ao* encoded a protein that localized to the core histone gene promoters. Moreover, it demonstrated that *ao^1^*/*ao^2^ trans*-heterozygous females produce eggs with significantly increased histone expression levels, and that reducing the histone gene copy number in *ao^1^*-homozygous females partially ameliorated the *ao-*associated maternal-effect lethality (Berloco *et al*. 2001). Together, these findings led to the conclusion that excess histone production in *ao* mutants caused the maternal-effect lethal phenotype. These results also suggested that heterochromatin could act as a ‘sink’ for excess histones, explaining why excess AO heterochromatin could alleviate the embryonic lethality associated with the loss of maternal *ao*. Thus, this landmark study linked the function of *ao*, a euchromatic gene that apparently controlled histone gene expression, to heterochromatin content and, ultimately, embryonic viability.

In *D. melanogaster*, histone genes are arranged in a tandemly repeated, multigene array of approximately 100 units, each comprising all four core histones (H2A, H2B, H3, and H4) and the linker histone (H1) on the 2^nd^ chromosome. Thus, on average, a diploid *D. melanogaster* genome encodes 200 such units, even though histone copy number varies significantly within *D. melanogaster* strains and between *Drosophila* species (Lifton *et al*. 1978; Strausbaugh and Weinberg 1982; Kremer and Hennig 1990; Mckay *et al*. 2015; Shukla *et al*. 2024). Although recently developed transgenic tools allow for more facile manipulation of histone gene copy numbers *in vivo* (Cook *et al*. 2012; Mckay *et al*. 2015; Zhang *et al*. 2019), the tight regulation of histone expression still makes manipulating histone expression levels in *Drosophila* challenging. For example, flies carrying 24 copies of the histone genes have nearly identical levels of histone transcripts and proteins as wild-type flies carrying 200 copies of the histone genes, likely due to a feedback-based compensation mechanism that ensures adequate histone expression levels regardless of histone gene copy number (Mckay *et al*. 2015). Thus, in addition to its exciting biology, *ao* emerged as a promising tool for manipulating histone gene expression in *D. melanogaster* (Chari *et al*. 2019).

Existing *ao* reagents, however, carry several caveats. First, only the *ao^1^*strain is still available, whereas the *ao**^2^*** strain has been lost. Second, the two mutants exhibit different phenotypes: *ao**^1^*** is viable as a homozygous mutant, but *ao**^2^*** was reported to be lethal as a homozygote (Tomkiel *et al*. 1995). Furthermore, although both *ao^1^/ao^1^* and *ao^1^/ao^2^* mutants exhibited maternal-effect lethality, more embryos from *ao^1^/ao^1^* mothers died at earlier stages compared to those from *ao^1^/ao^2^* mothers (Tomkiel *et al*. 1995). Finally, *ao^1^/ao^1^*stocks are unstable and can rapidly acquire genetic suppressors that alleviate the maternal-effect lethality (Krider and Levine 1975; Graziani *et al*. 1981; Manzi *et al*. 1986). These observations implied that genetic background effects could dramatically affect the severity of the phenotype associated with loss of *ao*.

To overcome these hurdles and to accurately characterize the *ao* phenotype, we used CRISPR/Cas9-based methods to generate new *ao* reagents: a precise knockout of *ao* to enable genetic analyses and V5 epitope*-*tagged alleles of *ao* at the endogenous locus to enable cytological visualization. Using these reagents, we recapitulated several classical genetic attributes of *ao*, including its maternal-effect lethality, which is suppressed either by a reduction in histone gene copy number or by excess heterochromatin on the Y chromosome. However, we could not confirm previous reports that Ao localizes to the histone gene cluster. We also discovered that *ao* does not affect histone transcript levels. Unlike in *ao^1^/ao^2^* flies, histone levels are unaffected in ovaries from both Δ*ao/*Δ*ao* and *ao^1^/ao^1^* homozygous females. Thus, although *ao* genetically interacts with heterochromatin as proposed in Sandler’s original hypothesis, we conclude that the molecular basis for these interactions still remains undiscovered.

## Materials and Methods

### Generation of the Δ*ao* line

We used CRISPR/Cas9 to create an *ao* knockout line. To facilitate the screening of transgenic flies, we replaced the *ao* allele with *dsRed* under the *3xP3* eye-specific promoter. We chose guide RNAs with the best efficiency score and no predicted off-targets (https://www.flyrnai.org/crispr/). We cloned guide RNAs (AGCCGGGTTCTTCTTCCGAT and AGTAATGTCTTTATTTACAA) targeting the 5’ and 3’ ends of the *ao* gene into pCFD4 U6:1_U6:3tandemgRNAs (Port *et al*. 2014) (Addgene plasmid #49411). The repair template sequence, including homology arms spanning approximately 1kb upstream and downstream of the *ao* coding sequence, was cloned into pDsRed-attP (Gratz *et al*. 2014) (Addgene plasmid #51019). To prevent guide RNAs from targeting PAM sites, we mutated the PAM sites on the repair template using the Q5 Site-Directed Mutagenesis Kit (New England Biolabs).

BestGene Inc. (Chino Hills, CA) prepared and co-injected the plasmids into BDSC 51323 (Bloomington Drosophila Stock Center) embryos expressing *vas-Cas9* on the X chromosome. Following injection, BestGene Inc. crossed the injected flies to a *yw* strain to isolate transformants, crossed out the *Cas9* gene, and balanced the 2^nd^ chromosome over *CyO.* We verified the absence of *ao* and the presence of *dsRed* with PCR and Sanger sequencing (see Table S1 for primer sequences). We extracted genomic DNA with the DNeasy Blood & Tissue Kit (Qiagen) according to the manufacturer’s protocol for insect tissue, then performed PCR using the Platinum PCR SuperMix High Fidelity (Invitrogen). The penetrance of the *CyO* phenotype decreased with temperature, which made it difficult to distinguish between homozygote-null and heterozygous flies. We, therefore, rebalanced the Δ*ao* allele over *CyO-gfp* marked with *mini-white*, which enabled us to screen for homozygous flies based on eye pigment color. To allow for *mini-white* visualization, the *CyO-gfp* strain carries *yw* on the X chromosome. We crossed the *yw* strain from BestGene Inc. with our *yw; CyO-gfp* strain to obtain a near-isogenic strain to our Δ*ao* strain and used this strain as the wildtype control for all experiments with Δ*ao* flies.

To obtain a Δ*ao* strain with a heterozygous deletion of the histone gene cluster, we crossed our Δ*ao* flies into the BDSC 8670 strain (Cook *et al*. 2012), which has a heterozygous deletion on the 2^nd^ chromosome corresponding to the histone gene array (chromosomal locus 39D3 to 39E2). Both the *ao* allele and histone genes are located on the left arm of the 2^nd^ chromosome. So, after obtaining a female fly heterozygous for both Δ*ao* and histone deficiency, we relied on recombination to obtain a fly with Δ*ao* and histone deficiency on the same 2^nd^ chromosome. We used this fly to make the Δ*ao, his(df)* stock. The histone deletion is not marked, so we used PCR using the Phusion High-Fidelity DNA polymerase (New England Biolabs) to determine its presence in the founder fly (see Table S1 for primer sequences).

### DNA isolation and Sanger sequencing

To confirm the endogenous Δ*ao* and *dsRed* insertion, we collected homozygous adult transgenic Δ*ao* flies from the heterozygous balanced stock (*y-, w-*; Δ*ao, 3xP3-dsRed/CyO, twi-GAL4, UAS-GFP*) and wild-type (*y-, w-*) flies. To isolate the DNA, we collected flies in 1.5mL microcentrifuge tubes and euthanized them at −80°C for 10 minutes. We added 50μL of squish buffer (10 mM Tris-HCl pH 8.0, 1 mM EDTA, 25 mM NaCl, and 200 μg/ml Proteinase K, with the enzyme added fresh from −20°C storage) to each tube, finely grinding the samples with p200 micropipette tips. We incubated the samples at 37°C for 30 minutes, then at 95°C for 2 minutes in heating blocks. We spun the samples at 20,000 RCF (relative centrifugal force) for 7 minutes and transferred the supernatant containing genomic DNA to new microcentrifuge tubes. We stored the samples overnight at 4°C. We performed 50μL PCR reactions using Q5 High-Fidelity 2X Master Mix (New England Biolabs) with primers designed to anneal upstream and downstream of the insertion (see R2 and F1 in Table S1 for primer sequences). We purified the PCR product with the Monarch PCR & DNA Cleanup Kit (New England Biolabs). We designed primers (Integrated DNA Technologies) to tile both the wild-type and Δ*ao* allele locus. Lastly, we performed Sanger sequencing (Azenta; primers listed in Table S1) and aligned the sequencing data to the endogenous *ao* and Δ*ao* sequences using SnapGene 5.1.7.

### Generation of the *ao-*transgene rescue line

We used the PhiC31 integrase system to create the *“*rescue” line with wild-type *ao*. We cloned the *ao* coding sequence (lacking the intron) with its endogenous promoter into the pattB plasmid, which contains an *attB* site and *mini-white* marker (Bischof *et al*. 2013)(DGRC stock 1420). BestGene Inc. prepared and injected the plasmid into BDSC 9750 embryos that carry the VK33 *attP* landing site on the 3^rd^ chromosome (Venken *et al*. 2006). BestGene Inc. confirmed the successful integration with PCR to verify the presence of the recombined *attL* site and the absence of the original *attP* site (see Table S1 for primer sequences). Then, the gene encoding the PhiC31 integrase was crossed out, and the 3^rd^ chromosome was balanced over *TM6B*.

### Generation of the V5-tagged *ao* line

We used CRISPR/Cas9 to tag *ao* at its C-terminus with a V5 epitope. We chose a guide RNA closest to the stop codon of *ao* with no off-targets (http://targetfinder.flycrispr.neuro.brown.edu/). We cloned the guide RNA (GTATAACCACAGCACAATAG) into pCFD5 (Port and Bullock 2016)(Addgene plasmid #73914). We designed a single-stranded oligonucleotide donor (ssODN) repair template containing the V5 tag (42 bp) and approximately 55 bp of sequence upstream and downstream of the insertion site. The ssODN had a mutated PAM site to prevent re-targeting.

We sent the midi-prepped (Qiagen) pCFD5 plasmid containing the guide RNA and lyophilized ssODN to GenetiVision Inc. (Houston, TX) for injection into embryos expressing *nanos*-*Cas9.* We screened for transformants using a PCR strategy, with primers that annealed upstream and downstream of the insertion site (see Table S1 for primer sequences). We tested for insertion of the V5 tag by the presence of a 42 bp shift in band size. Finally, we confirmed the successful insertion of the intact V5 tag by Sanger sequencing.

### Fly husbandry, fertility, and viability assays

We maintained flies on the benchtop at room temperature on corn syrup/soy media made in-house at Fred Hutchinson Cancer Center (Seattle, WA) or purchased from Archon Scientific (Durham, NC). To conduct fertility assays, we used 1-to-5-day-old males and virgin females raised at room temperature. Unless otherwise noted, we paired four virgin females with two males in a vial with corn syrup/soy media and allowed them to mate for three days (for X^Y assays) or one week (for all other assays). To prevent larval overcrowding in the vials, we flipped the parents to new vials after three days and discarded the parents from the new vials four days later. As noted for each experiment, we set up and maintained the crosses at 18°C, 25°C, or 29°C. We counted the adult offspring (F1) until no more progeny were produced. Data for all fertility and viability assays are available in Table S2.

We excluded crosses with no larvae in either vial from statistical analyses. Because non-genetic factors can influence fly fertility, including variation in food and ambient humidity, we compared data only within crosses set up on the same day. We used GraphPad Prism version 10.1.1 for macOS (GraphPad Software) to plot the data and conduct statistical analyses. We performed two-tailed Mann-Whitney U tests to compare the offspring counts between the two datasets and reported the exact *p*-values. To compare the observed offspring genotype to the theoretical Mendelian offspring genotype, we used a one-sample proportion test (http://www2.psych.purdue.edu/~gfrancis/calculators/proportion_test_one_sample.shtml). To compare the results of X^Y crosses, we analyzed a 2×2 contingency table using a two-tailed Fisher’s exact test (https://www.graphpad.com/quickcalcs/contingency1/).

We performed late-stage developmental viability assays as previously described (Spring *et al*. 2019). Briefly, we transferred 40-50 second-instar larvae into vials containing standard molasses fly food and waited for them to complete development. We counted the number of pupal cases in each vial and the number of eclosed adults. We calculated the percentage viability at the pupal and adult stages by dividing each value by the initial number of larvae and multiplying by 100. Each vial represents a single biological replicate, and we had 5-8 replicates for each genotype.

### Immunofluorescence

#### Ovaries

The tagged V5-Lsm11 control stock (V5-Lsm11 pAttB10A, Lsm11^c02047^/CyO, *twi*-GAL4, UAS::GFP) was a gift from Dr. Robert Duronio (Godfrey *et al*. 2009). We dissected ovaries in PBS, then fixed with 1:1 paraPBT: heptane (paraPBT = 4% paraformaldehyde in PBS + 0.1% Triton X-100) in an Eppendorf tube for 10 minutes at room temperature. Following three 5-minute washes in PBST (PBS + 0.1% Triton X-100), we blocked the ovaries in PBST with 3% BSA for 30 minutes at room temperature. We incubated the ovaries in primary antibodies overnight at 4°C. We used a guinea pig anti-Mxc antibody (gift from Dr. Robert Duronio)(White *et al*. 2011) at 1:5000 and an anti-V5 Tag monoclonal antibody (Thermo R960-25) at 1:250. After three 5-minute washes in PBST, we incubated the samples with secondary antibodies in PBST for 2 hours at room temperature. We used the goat anti-mouse IgG Alexa Fluor 568 (Thermo A-11031) and goat anti-guinea pig IgG Alexa Fluor 488 (Thermo A-11073), both at 1:2000. We added Hoechst stain (Invitrogen) to the samples in the last 30 minutes of the incubation with the secondary antibody. After three 5-minute washes with PBST, we mounted the ovaries onto slides with 20 μL of SlowFade Gold Antifade Mountant with DAPI (Invitrogen), then added coverslips and sealed with nail polish.

#### Polytene chromosomes

To overexpress *ao* in larval salivary glands, we collected virgin w[1118]; P(w[+mC]=Sgs3-GAL4.PD)TP1 (Bloomington Stock Center #6870) females and bred them to either male *y*-, *w*- flies as a control or to male (UAS- ao.ORF.3xHA.GW)ZH-86Fb flies (Fly ORF #6093) (Bischof *et al*. 2013) at 23°C. Chromosomes from F1 or from V5-tagged *ao* lines were prepared as previously described (Hodkinson *et al*. 2024). We dissected salivary glands from 3rd-instar larvae and fixed them after three washes. First, we used a 4% paraformaldehyde and 1% Triton X-100 solution in 1X phosphate-buffered saline (PBS) for 1 minute. We then transferred the glands to a 4% paraformaldehyde and 3M acetic acid solution for two minutes. Finally, we transferred the glands to a solution of 1 part lactic acid, 2 parts H2O, and 3 parts acetic acid for 5 minutes. We placed fixed glands on a coverslip and crushed them with a slide to spread the polytene chromosomes. We plunged the slides in liquid N2 and removed the coverslips. We stored the slides in 95% ethanol at - 20°C. Before staining, we rehydrated the slides in 1X PBS for 15 minutes, then permeabilized them in 1% Triton X-100 for 10 minutes. We blocked the glands in 0.5% BSA in 1X PBS for one hour at room temperature. The 0.5% BSA solution was also the diluent for the antibodies. We dispensed the primary antibody solution onto a coverslip on the slide and incubated it overnight at 4°C. The following day, we washed off the primary antibody solution with three 15-minute washes in 1X PBS. Likewise, we dispensed a secondary antibody solution and placed a coverslip over the sample. We incubated these in the dark at room temperature for two hours. Following incubation, we washed the slides three times for 10 minutes in 1X PBS, dried them, mounted them with ProLong Diamond Anti-Fade Mountant with DAPI (P36962, Thermo Fisher Scientific), sealed the coverslip to the slide with nail polish, and stored them at 4°C. We imaged polytene chromosomes with a ZEISS Axio Scope.A1 fluorescence microscope and the others with a Keyence BZ-X810 All-in-One-Fluorescence Microscope using the “BZ-X800 Viewer” program.

### RNA extraction and quantitative reverse transcriptase PCR (RT-qPCR)

We dissected Δ*ao* or *ao^1^* ovaries from 4-day-old virgin females in PBS. To collect unfertilized eggs, we allowed 3- to 7-day-old virgin females to lay on grape plates for 7 hours. We transferred 4 pairs of ovaries, or 10 unfertilized eggs, into an Eppendorf tube. We homogenized the tissue in 20 μL of TRIzol (Invitrogen) using a disposable pestle and an electric homogenizer. We stored samples at −80°C in 100 μL of TRIzol until ready for processing. We incubated the thawed samples in 1 mL of TRIzol for 5 minutes, then centrifuged them at 13,000 rpm for 10 minutes at 4°C to separate the supernatant. We extracted the supernatant with chloroform and extracted the soluble phase with isopropanol. After a wash in 70% ethanol, we resuspended the RNA pellet in RNase-free water. We treated the samples with DNase I (Zymo Research) and then purified them with the RNA Clean & Concentrator-5 kit (Zymo Research). We quantified the purified samples using the Qubit RNA Broad Range Assay Kit (Invitrogen), then synthesized cDNA with the SuperScript III First-Strand Synthesis System using random hexamers (Invitrogen).

To perform RT-qPCR on Δ*ao* or *ao^1^* ovaries and Δ*ao* unfertilized eggs (Fig. 3a, Fig. 3b, Fig. S9, Fig. S15a), we used the PowerUp SYBR Green Master Mix for qPCR (Applied Biosystems) with approximately 10ng of cDNA per reaction. We used the QuantStudio 3 Real-Time PCR System (Applied Biosystems) to run the RT-qPCR experiment using the *rp49* gene as a control (see Table S1 for primer sequences). We ran each sample in technical triplicate for each primer pair and used the median value for analysis.

To measure histone transcript levels in unfertilized eggs from *ao^1^/ao^1^* homozygous mothers (Fig. S15b), we used a different set of primers (see Table S1) and AzuraQuant Green Fast qPCR Mix LoRox (Azura Genomics). We report a mean of 2 technical duplicates and 4-5 biological replicates. We used the same setup to measure *ao* transcript levels in ovaries from 3-day-old virgin *ao-V5* flies (Fig. S13) and in ovaries from *D. melanogaster* strains with different histone copy numbers (Fig. 4b).

To measure *ao* mRNA levels after GAL4-UAS overexpression in salivary glands (Fig. S12), we isolated RNA from dissected salivary glands of 3rd instar larvae (10 pairs per replicate), placed them in 100 μL TRIzol, homogenized with plastic pestles, and stored them at −80°C. We later brought samples to 1mL in Trizol and rotated them for 20 min. Subsequently, we added 200 μL chloroform to the tubes and shook them vigorously, then spun them down at 20,000 RCF for 20 minutes at 4°C. We transferred the aqueous phase to a new tube and repeated the chloroform addition, shaking, spinning, and aqueous-phase removal. We added 500 μL isopropanol and 5μL glycogen to the final aqueous layer, then incubated the samples at −20°C overnight to precipitate RNA. We spun the samples at 20,000 RCF and 4°C for 10 minutes, then washed the RNA pellets with 75% ethanol. We spun the samples again in a centrifuge for 5 minutes at 20,000 RCF. Finally, we resuspended RNA in water and stored it at −80°C until use. For RT-qPCR, we utilized a LunaScript RNA-to-cDNA conversion kit (NEB) to prepare samples. We used the RT-qPCR primers for *ao*, *ao*-HA, and *rp49* in Table S1. We report a mean of 2 technical duplicates and 1-5 biological replicates. Biological replicates in this experiment were produced from different crosses performed asynchronously. We performed qPCR from biological replicates on separate occasions.

To analyze the RT-qPCR data, we normalized gene expression to the reference gene *rp49* and calculated fold change using the 2^−ΔΔCt^ method (Livak and Schmittgen 2001). For Fig. 3a, Fig. 3b, Fig. 4b, Fig. S9, and Fig. S15, we calculated ΔΔCt by subtracting the ΔCt of the paired control sample from that of the experimental sample, and used the resulting 2^−ΔΔCt^ value (fold change). We then used the one-sample t-test to compare the fold change in gene expression (normalized to *rp49* expression) of the experimental genotype relative to the control genotype.

For Fig. S12 and Fig. S13, we first calculated the average 2^-ΔCt^ for the control genotype. We then calculated the normalized relative gene expression for each experimental sample by dividing the 2^-ΔCt^ of the sample by the average 2^-ΔCt^ for the control genotype. For experiments run on more than one plate (Fig. S12), we compared the 2^-ΔCt^ to the average 2^-ΔCt^ for the control genotype on the same plate. We then used the Welch’s t-test to compare the fold change in *ao* expression (normalized to *rp49* expression) of each experimental genotype relative to the control genotype.

### Sample preparation and data analysis for the ribo-depletion RNA-seq analyses

We dissected ovaries from 4-day-old Δ*ao* and isogenic *yw* virgin females for RNA-seq. We prepared biological triplicate samples for each genotype, with four pairs of ovaries per replicate. Each Δ*ao* replicate sample was paired with an isogenic *yw* sample, with each paired sample set collected on the same day to account for environmental factors that might affect gene expression (*e.g.,* changes in humidity, temperature, or food composition). We transferred four pairs of ovaries to an Eppendorf tube for each replicate and homogenized the tissue in 20 μL of TRIzol (Invitrogen) using a disposable pestle and an electric homogenizer. We then stored the samples at −80°C in 100 μL of TRIzol until they were ready for processing. We processed all six samples in parallel on the same day to ensure minimal variation during processing. We incubated the thawed samples in 1 mL of TRIzol for 5 minutes, then centrifuged at 13,000 rpm for 10 minutes at 4°C to separate the supernatant. We extracted the supernatant with chloroform, then the soluble phase with isopropanol. After washing the RNA pellet with 70% ethanol, we resuspended it in RNase-free water. We treated the samples with DNase I (Zymo Research) and purified them with the RNA Clean & Concentrator-5 kit (Zymo Research). We quantified the purified samples with the Qubit RNA Broad Range Assay Kit (Invitrogen), then sent the RNA to Azenta Life Sciences (Burlington, MA) for ribosomal RNA depletion library preparation and sequencing. Azenta added ERCC (External RNA Controls Consortium) RNA Spike-In Controls mixes to the samples before library preparation.

We mapped the reads to the *D. melanogaster* dm6 genome assembly using HISAT2 (Kim *et al*. 2019), which aligns repetitive reads to a single genomic location. We utilized the multicov subfunction in the BEDTools suite (Quinlan and Hall 2010) to count histone reads, and used R to sum the counts at all loci for each histone gene. We combined the summed histone counts with single-copy gene counts and used DESeq2 (Love *et al*. 2014) to detect differential gene expression between Δ*ao* and wild-type samples. Data analyses were performed using R 4.2.2 (R Core Team, 2022) and Galaxy (The Galaxy Community, 2024).

### Western blotting

We dissected ovaries from four-day-old virgin females in PBS on ice. We transferred five pairs of ovaries to an Eppendorf tube, added 20 μL of 2x Laemmli Buffer (Bio-Rad Laboratories) with 200 mM DTT and 300 mM NaCl, and flash-froze the samples in liquid nitrogen. The samples were stored at - 80°C until processing. To extract protein, we thawed the samples on ice and added protease inhibitors (EDTA-free cOmplete ULTRA Tablets, Roche). Then, we hand-pestled the samples on ice using disposable pestles. After a brief spin at 4°C to collect the samples, we boiled them at 100°C for 10 minutes, then centrifuged them at maximum speed for 2 minutes to obtain a clean lysate.

We loaded 10 μL of protein sample per well on the Any kD Mini-PROTEAN TGX Precast Protein Gel (Bio-Rad Laboratories). We ran the gel for 90 minutes at 100 V in Tris/Glycine/SDS buffer, then transferred the gel to a Trans-Blot Turbo Mini 0.2 μm Nitrocellulose membrane (Bio-Rad Laboratories) using the Trans-Blot Turbo Transfer System (Bio-Rad Laboratories). We washed the membrane three times with PBS, blocked in Intercept (PBS) Blocking Buffer (LI-COR) for 1 hour at room temperature, then probed with primary antibodies in phosphate-buffered saline with 0.1% Tween-20 (PBST) at 4°C overnight. We used the following primary antibodies: rabbit anti-beta-tubulin (abcam 6046) at 1:1000, mouse anti-H2B (abcam 52484) at 1:3000, and rabbit anti-H3 (abcam 1971) at 1:4000. Following three 10-minute washes in PBST, we incubated the membrane for 1 hour at room temperature with IRDye 680RD Donkey anti-Mouse IgG (LI-COR) and/or IRDye 800CW Donkey anti-Rabbit IgG 800 (LI-COR) secondary antibodies in PBST at 1:20,000. After three washes with PBST and a final wash in PBS, we scanned the membrane at 700 nm and 800 nm on the Odyssey CLx Imager (LI-COR). We quantified the blots using Image Studio v6.0 (LI-COR). We used the Manual analysis option with median background correction (border 3 for all segments). For each pair of samples, we used the “Add Rectangle” function to draw a box around the larger band, then added a box of the same size to the other band using the “Add Selection” function. We normalized H2B or H3 expression to beta-tubulin expression as a loading control, then normalized the Δ*ao* sample to the corresponding wildtype (*yw*) sample to obtain relative quantification.

## Results

### Δ*ao-*knockout flies have partial maternal-effect lethality

To obtain an *ao* mutant without genetic background effects, we used CRISPR/Cas9 to create a Δ*ao* strain using guide RNAs designed to target the start of the 5’ UTR and the end of the 3’ UTR of *ao* (Fig. 1a). To facilitate the phenotypic screening of Δ*ao* flies, we inserted a repair template with the fluorescent marker *dsRed* under the control of the eye-specific promoter *3xP3* using homology arms of approximately 1000 base pairs. We verified the *ao* knockout and *dsRed* replacement, as well as the integrity of the upstream gene *ATPsynG*, using PCR and Sanger sequencing (Fig. S1-S2). We also generated a nearly isogenic strain to the *yw;* Δ*ao/CyO-gfp* strain except for a wild-type 2^nd^ chromosome in place of Δ*ao*, which we used as the ‘wild-type’ control for all future experiments.

**Figure 1.**
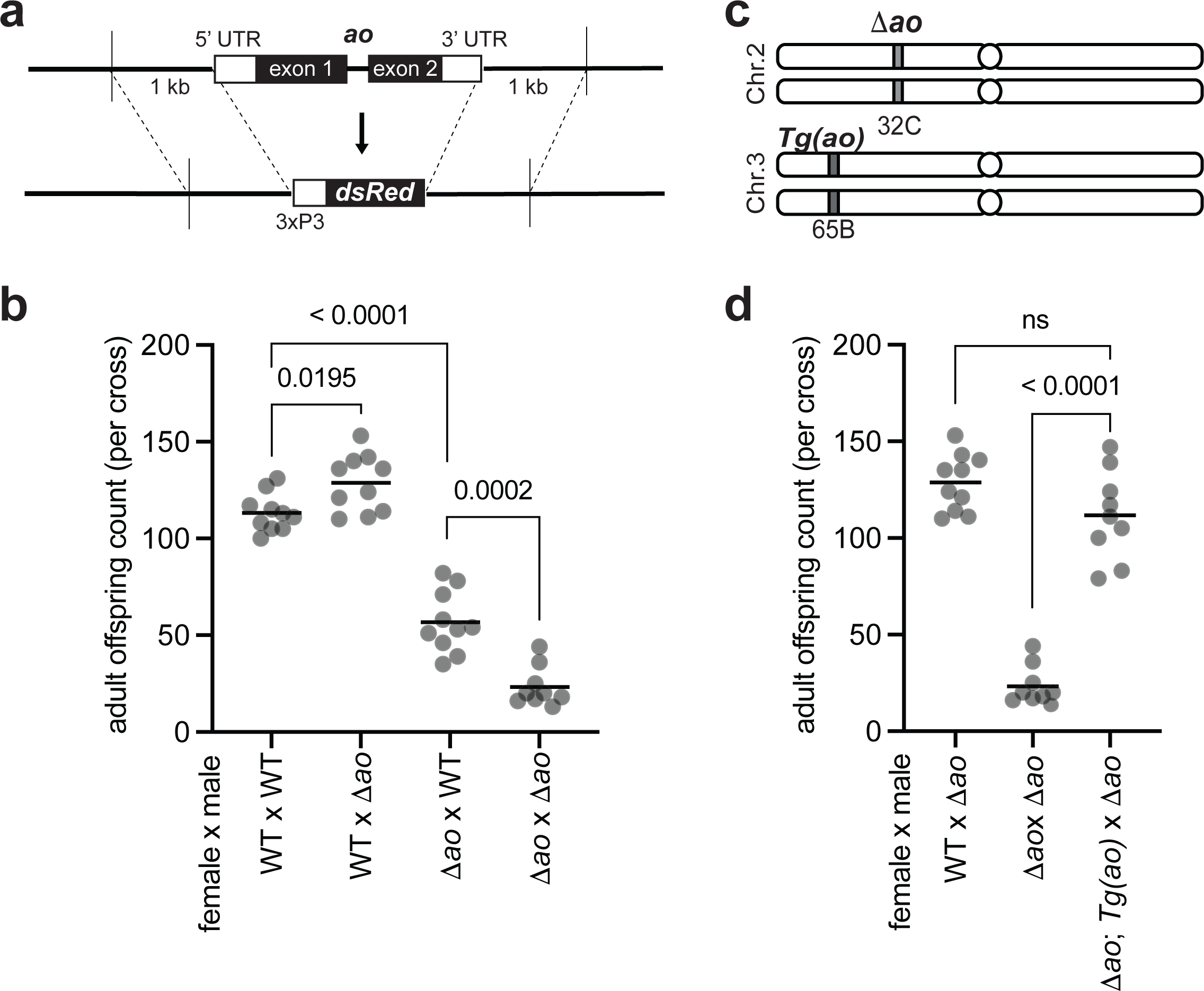
Loss of *ao* causes partial maternal-effect lethality in *D. melanogaster.* **(a)** We replaced the *ao* coding sequence and UTRs with *dsRed* (fluorescent marker) under the control of *3xP3*, an eye-specific promoter, using CRISPR-Cas9 and homology arms (detailed in Fig. S1). **(b)** Crosses between Δ*ao* females and wildtype males yield fewer adult progeny than crosses between wildtype females and wildtype males or between wildtype females and Δ*ao* males, confirming that loss of *ao* leads to partial maternal-effect lethality, which is further exacerbated in crosses between Δ*ao* females and Δ*ao* males. The mild increase in offspring number in the cross between wildtype females and Δ*ao* males is not reproducible (Fig. S3). Each point on a graph is the offspring count from a biological replicate cross performed at 29°C. The *p-*values from two-tailed Mann-Whitney U tests are shown above the compared samples. **(c)** We integrated a ‘rescue’ *ao* transgene construct on the *D. melanogaster* 3^rd^ chromosome using the PhiC31 integrase system (details in Fig. S8). The ‘rescue’ *ao* transgene is expressed at ∼20% of the levels of the endogenous *ao* gene (Fig. S9). **(d)** Despite its lower expression, the ‘rescue’ *ao* transgene can restore the number of adult offspring produced from Δ*ao* females to nearly wildtype levels at 29°C. The *p-*values are from two-tailed Mann-Whitney U tests.

We first assessed whether our newly generated Δ*ao* strain recapitulated the maternal-effect lethality phenotype of *ao* mutants (Sandler 1970; Sandler 1977; Tomkiel *et al*. 1995). Since *ao*-associated maternal-effect lethality was more pronounced at higher temperatures (Sandler 1970), we measured the total number of adult offspring produced by Δ*ao* flies at 29°C. Consistent with previous findings, we found that Δ*ao* females exhibit partial maternal-effect lethality when crossed to wild-type males (Fig. 1b). We found that maternal-effect lethality is exacerbated in crosses between Δ*ao* females and Δ*ao* males (Fig. 1b), confirming previous findings that a paternal copy of the wild-type *ao* allele can partially rescue zygotic survival (Pimpinelli *et al*. 1985; Tomkiel *et al*. 1995). Although our data initially suggested a slight fertility increase of Δ*ao* males relative to wildtype males in crosses with wildtype females (Fig. 1b), subsequent experiments revealed no significant differences in these crosses (Fig. S3).

We confirmed our findings by measuring the survival of pupae or adults from a given number of larvae produced from *ao^1^/ao^1^* mutant mothers (Fig. S4). This assay examines viability at later developmental stages rather than across all stages (Fig. 1b). Nevertheless, these findings (Fig. S4) were nearly identical to our findings of offspring viability from Δ*ao/*Δ*ao* mothers (Fig. 1b). Thus, the maternal-effect lethality resulting from *ao* loss continues to manifest at both early and later stages of development. It can be further exacerbated by the loss of a paternal (and zygotic) *ao*.

We further assessed zygotic effects, as measured by survival of Δ*ao* flies, by looking for deviations from the expected Mendelian ratio in offspring genotypes. Without a zygotic effect, the theoretical Mendelian ratio for offspring from two heterozygous parents should be 33% Δ*ao* homozygotes and 66% heterozygotes (since homozygosity for balancer chromosomes leads to lethality). In contrast to this expectation, the observed offspring genotype ratio from parents heterozygous for Δ*ao* was 28% homozygous and 72% heterozygous, indicating a mild but statistically significant zygotic effect (one-sample proportion test, *p =* 0.0043; Fig. S5).

We found that heterozygous females with one copy of the Δ*ao* allele produce the same number of adult offspring as wild-type females when crossed to Δ*ao/*Δ*ao* males (Fig. S6) (Sandler 1970). Finally, we found that *ao*-associated maternal-effect lethality is most pronounced at 29°C but also manifests at 25°C and 18°C, albeit to slightly lower extents (Fig. S7), consistent with previous findings (Sandler 1970). Thus, the Δ*ao* strains we created confirm previous findings from *ao^1^ and ao^2^* strains: loss of *ao* results in no significant consequences to paternal fertility but does cause maternal-effect lethality, which can be partially rescued by a wild-type paternal allele of *ao* in the zygote.

We generated a ‘rescue’ *ao* transgene to rule out the possibility that the maternal-effect lethality in Δ*ao* strains might have resulted from an off-target mutation introduced during the CRISPR/Cas9 cleavage or repair (Fig. 1c). We used the PhiC31 integrase system (Groth *et al*. 2004) to insert an *ao* transgene onto the 3^rd^ chromosome, flanked by approximately 700 base pairs upstream and 300 base pairs downstream of the *ao* protein-coding sequence. Since the *ao* promoter is poorly defined, we included the untranslated regions of neighboring genes but not their coding sequences. By crossing the Δ*ao* and ‘rescue’ *ao* strains, we obtained flies with both homozygous Δ*ao* mutations (on the 2^nd^ chromosome) and two copies of the ‘rescue’ *ao* transgene (on the 3^rd^ chromosome) (Fig. S8). The ‘rescue’ *ao* transgene is transcribed at only 20% of the level of the endogenous *ao* gene in ovaries (Fig. S9, Table S3). Despite its lower expression, the ‘rescue’ *ao* transgene is sufficient to entirely suppress the maternal-effect lethality of Δ*ao* (Fig. 1d), thereby demonstrating a causal association of the maternal-effect lethality we observed with loss of *ao*.

### Ao does not localize to the histone gene cluster

Using a polyclonal antibody raised against the Ao protein, a previous study reported that Ao localizes to the multigene array of histone genes on the 2^nd^ chromosome in polytene chromosomes from salivary glands and in mitotic chromosomes from larval neuroblasts (Berloco *et al*. 2001). However, this antibody is no longer available. To visualize the Ao protein, we generated endogenously tagged *ao* alleles in which we tagged either the C-terminus (“*ao-V5*”) (Fig. 2a, Fig. S10) or N-terminus (“*V5- ao*”) (Fig. S11) of *ao* with a V5 epitope at the endogenous locus using CRISPR/Cas9. We confirmed that the *ao-V5* female flies have wild-type fertility at 29°C, indicating that the V5 tag does not interfere with Ao protein function (Fig. 2b).

**Figure 2.**
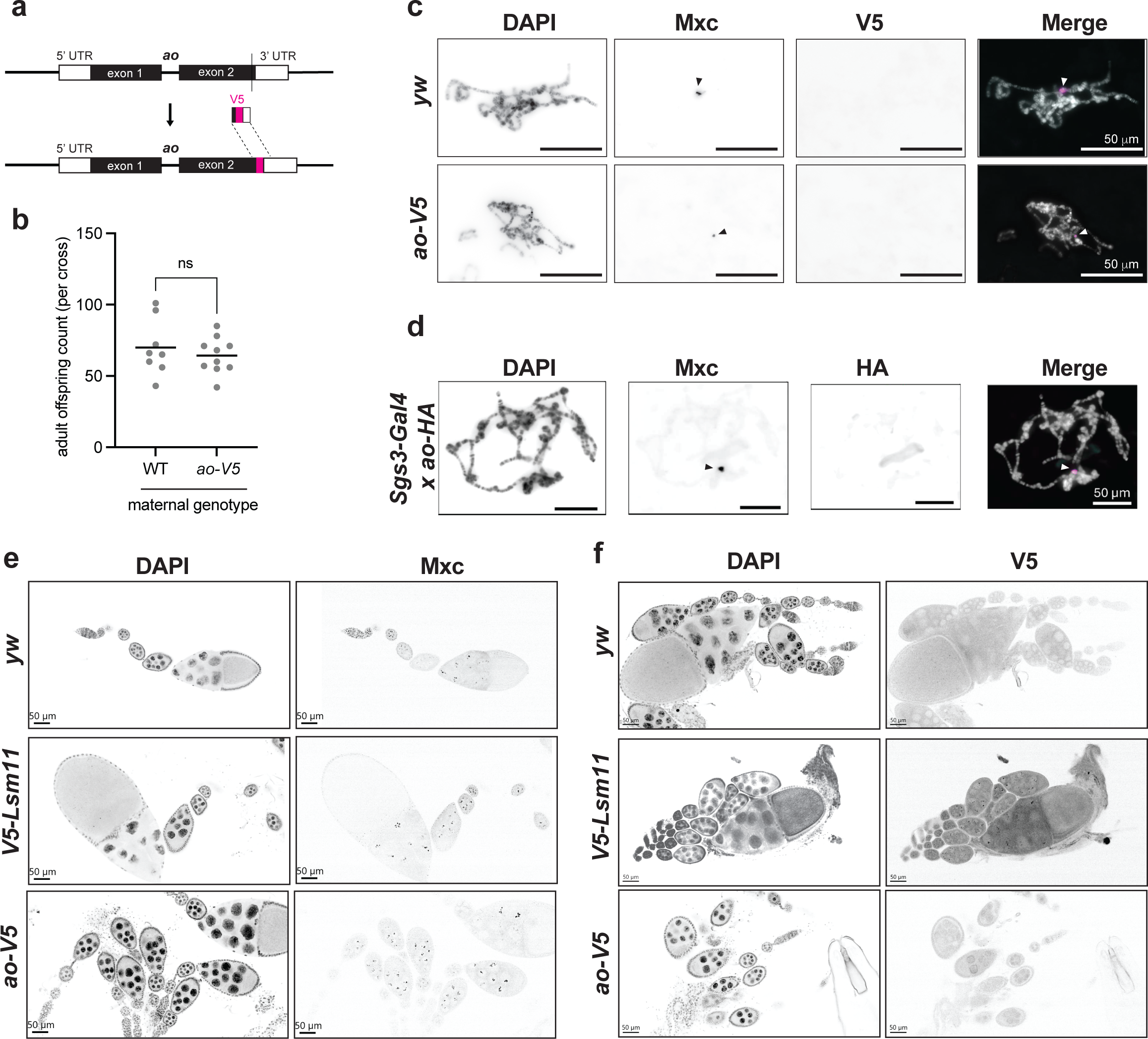
Ao-V5 does not localize to the histone gene locus in *D. melanogaster*. **(a)** Using CRISPR-Cas9, we introduced an in-frame V5 epitope tag at the 3’ end of the *ao* coding sequence at its endogenous locus (Fig. S10). **(b)** Crosses between *ao-V5* females and *yw* males yield the same number of adult offspring as those between *yw* females and *yw* males at 29°C. The *p-*values are from two-tailed Mann-Whitney U tests. **(c)** We performed immunofluorescence in polytene chromosomes from salivary glands dissected from *ao-V5* and *yw* (control) flies. We did not observe colocalization of Mxc (arrowhead), which localizes to histone locus bodies (HLBs), and Ao-V5. **(d)** We overexpressed *ao* in salivary glands using crosses between two *D. melanogaster* strains, one expressing a salivary gland-specific GAL4 driver (*Sgs3-GAL4*) and the other expressing an HA-tagged *ao* transgene (*UAS-ao-HA*). Despite dramatic *ao* overexpression in salivary glands (Fig. S12), we did not observe Ao-HA colocalizing with Mxc (arrowhead) on polytene chromosomes. **(e)** We found that Mxc forms puncta in ovarian nurse cells in all three *D. melanogaster* strains assayed: *yw, V5-Lsm11,* and *ao-V5*. **(f)** We assayed staining of the V5 epitope-tag in ovaries from three *D. melanogaster* strains: *yw* (which does not express a V5-tagged protein)*, V5-Lsm11* (which expresses V5-Lsm11, previously shown to localize to HLBs), and *ao-V5*. Expectedly, we observed Lsm11-V5 localizing in punctate dots in nurse cell nuclei (just like Mxc), but no V5 staining in either *yw* or *ao-V5* ovaries. In all cases, since we observed no discrete signal, we infer that epitope-tagged Ao lacks a discernible localization pattern on polytene chromosomes or in ovarian cells. (Scale bar = 50 μm).

Using the V5-tagged *ao* alleles, we first sought to confirm the observation by Berloco *et al*. (2001) that Ao localizes to the histone gene cluster on polytene chromosomes. We stained polytene chromosomes from salivary glands with antibodies against the V5 epitope and the Multi sex combs (Mxc) protein, which localizes specifically to the histone gene cluster and is a core structural component of the histone locus body (White *et al*. 2011). We did not observe colocalization of Ao-V5 and Mxc using the *ao-V5* allele (Fig. 2c) or the *V5-ao* allele (Fig. S11). Moreover, we found that Ao-V5 does not localize specifically to any chromosomal loci. Our inability to replicate prior observations (Berloco *et al*. 2001) could be because *ao* mRNA is nearly undetectable in salivary glands (Brown *et al*. 2014; Ozturk-Colak *et al*. 2024). We therefore overexpressed transgenic HA-tagged *ao* specifically in salivary glands using the GAL4-UAS system. Using this strategy, we observed a tremendous increase in *ao-HA* expression in salivary glands (Fig. S12, Table S4), yet Ao-HA still did not colocalize with Mxc at the histone genes in salivary glands (Fig. 2d), or to any distinct chromosomal loci.

Given *ao*’s maternal-effect lethal phenotype, we next investigated the expression and localization of the Ao-V5 protein in ovaries. To confirm that the V5 tag does not impair Ao expression, we first performed quantitative reverse-transcriptase PCR (RT-qPCR) for *ao* in ovaries. We found that *ao-*V5 homozygotes, but not heterozygotes, produce ∼1.5 times as much *ao* transcript as *yw* controls (Fig. S13, Table S5) without adversely affecting female fertility (Fig. 2b). We next stained ovaries for Ao-V5 and Mxc. Unfortunately, we found significant antibody cross-reactivity in this tissue, which prevented us from interpreting co-staining results. To overcome this potential artifact, we therefore separately stained for V5 and Mxc in three different strain backgrounds. The first is the ‘wild-type’ *yw* strain, which lacks Ao-V5 and serves as a negative control. The second is a *V5-Lsm11* strain (Godfrey *et al*. 2009), which serves as a positive control, since the Lsm11 protein targets the histone locus where it is involved in the 3’ end processing of histone transcripts (White *et al*. 2011). The third strain is the *ao-V5* strain we have constructed. Expectedly, we find that Mxc forms clear puncta in the ovarian nurse cells in all three strains (Fig. 2e). Also as expected, we found no V5 staining in the *yw* strain, but clear puncta in ovarian nurse cells in the *V5-Lsm11* strain (Fig. 2f). However, we found no V5 puncta in the *ao-V5* ovaries (Fig. 2f), even though this strain expresses higher than wild-type levels of *ao* (Fig. S13) and does not result in maternal-effect lethality (Fig. 2b). Based on our immunofluorescence experiments in multiple tissues with two different epitope tags, we cannot confirm prior reports of Ao localization to the histone gene cluster (Berloco *et al*. 2001). In addition, we found that Mxc foci are unaffected in Δ*ao/*Δ*ao* flies, suggesting that loss of Ao does not lead to disruption of histone locus bodies or Mxc localization (Fig. S14).

### *ao* does not affect histone transcript levels

The same previous study also reported the remarkable finding of significant overexpression of core histone genes in *ao* mutants (Berloco *et al*. 2001). Based on northern blotting analyses, the study noted that unfertilized eggs from *ao^1^/ao^2^* females had a 1.6-fold (for histone H4) to 11-fold (for histone H2A) increase in core histone transcript levels relative to unfertilized eggs from wild-type Oregon-R females (Berloco *et al*. 2001). To quantify histone transcript levels in ovaries and unfertilized eggs from virgin females from our Δ*ao* strain, we performed RT-qPCR for each of the four core histones and the linker histone, using a similar strategy as previously reported (Bulchand *et al*. 2010; Rieder *et al*. 2017) but with slightly different primers (see Methods, Table S1). Surprisingly, we found no significant differences in core histone levels between Δ*ao* and isogenic wild-type samples in both ovaries and unfertilized eggs (Fig. 3a, Table S6; Fig. S15a, Table S7).

**Figure 3.**
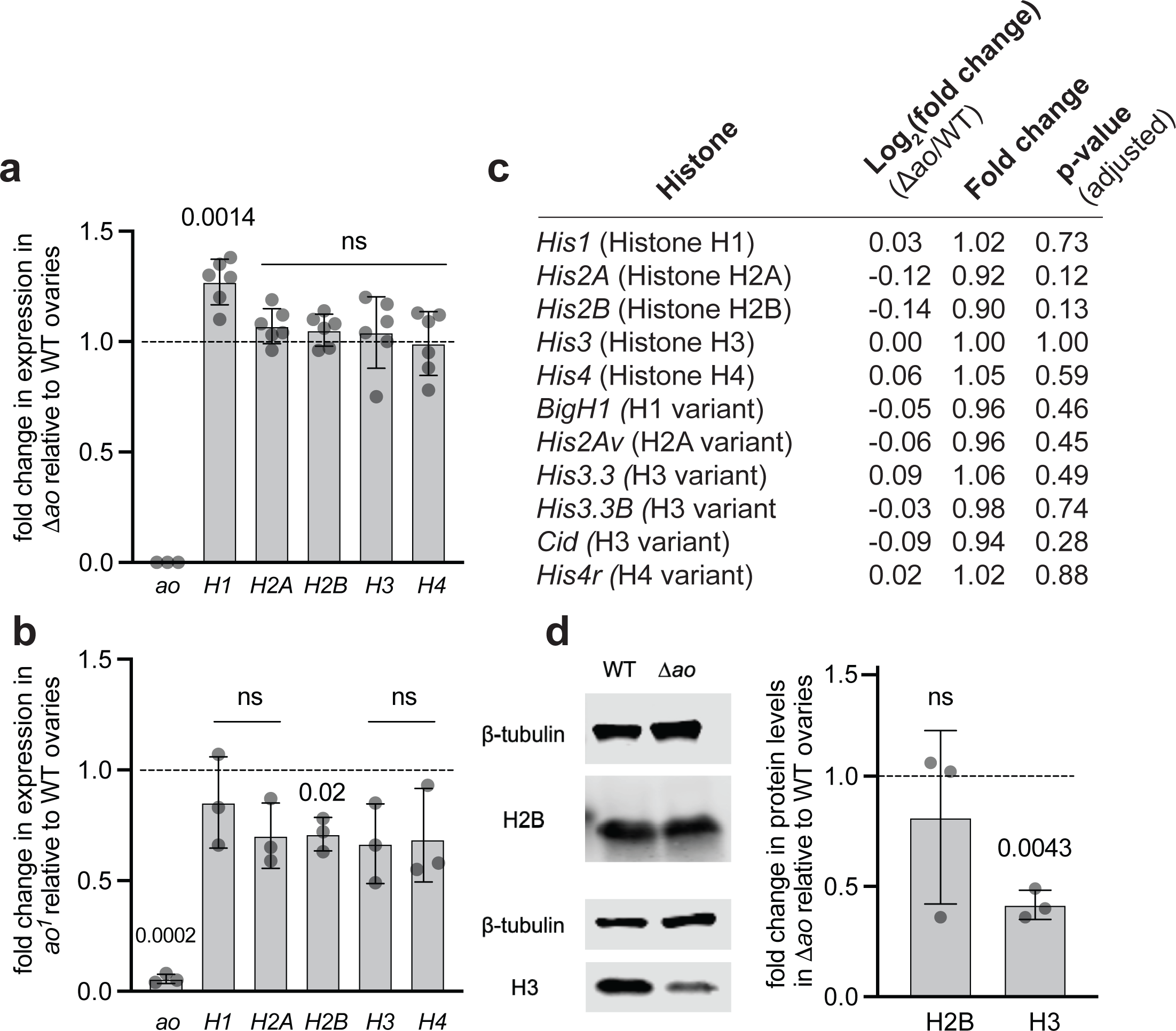
Histone expression levels in *ao* mutant ovaries. **(a)** RT-qPCR analyses measure the levels of the linker (H1) and core (H2A, H2B, H3, H4) histone transcripts in Δ*ao* ovaries relative to ovaries from isogenic *yw* females (dashed line). Each data point is a biological replicate of 3-5 pairs of ovaries from 4-day-old virgin females. For each replicate, the median of the technical triplicate is shown. Gene expression has been normalized to ribosomal protein *rp49.* The *p*-values are calculated using a one-sample t-test (Methods). The bars represent one standard deviation. **(b)** RT-qPCR on *ao^1^* ovaries presents the levels of the linker (H1) and core (H2A, H2B, H3, H4) histone transcripts relative to ovaries from *yw* females (dashed line). The *yw* strain is not isogenic to the *ao^1^* strain. Each data point is a biological replicate of 3-5 pairs of ovaries from 4-day-old virgin females. For each replicate, the median of the technical triplicate is shown. Gene expression has been normalized to *rp49.* The *p*-values are calculated using a one-sample t-test. The bars represent one standard deviation. **(c)** We performed RNA-seq on ovaries from Δ*ao* and isogenic *yw* 4-day-old virgin females. We used ribosomal RNA-depletion library preparation to capture replication dependent histones. Each genotype consisted of biological triplicates, with n=4 pairs of ovaries per replicate. Log_2_(fold change) represents the differential gene expression of Δ*ao* samples relative to the wildtype samples. The adjusted *p-*values were calculated using the False Discovery Rate/Benjamini-Hochberg procedure. **(d)** Western blots on Δ*ao* ovaries reveal no difference in H2B levels between Δ*ao* and wildtype (isogenic *yw*) ovaries, but 50% lower levels of H3 protein in Δ*ao* ovaries. We used beta-tubulin as a loading control for visualization and quantification. The *p*-values are calculated using a one-sample t-test. The bars represent one standard deviation.

Given the discrepancy between our Δ*ao/*Δ*ao* results and previous findings from *ao^1^/ao^2^* flies (Berloco *et al*. 2001), we further quantified histone transcript levels in *ao^1^/ao^1^* ovaries by RT-qPCR (the *ao^2^* stock no longer exists and cannot be assayed). Like in Δ*ao* ovaries, we found no evidence for a significant increase in core histone transcript levels in ovaries from *ao^1^/ao^1^* and wild-type females (Fig. 3b, Table S8). In contrast, we observed a mild but statistically significant *decrease* in core histone H2B transcripts (Fig. 3b, Table S8). Similarly, when assessing core histone transcript levels in unfertilized eggs from *ao^1^/ao^1^* females, we again observed either no difference or a slight decrease (for histone H2B) in core histone transcript levels (Fig. S15b, Table S9).

As an orthogonal method to confirm the results of the RT-qPCR assays, we also conducted RNA-sequencing to investigate whether the expression of any histone gene changed dramatically between Δ*ao* and wild-type ovaries. We used a ribosomal RNA-depletion library preparation to capture replication-dependent histones, which lack poly(A) tails (Marzluff *et al*. 2008). These analyses confirmed the results from our RT-qPCR analyses, showing that the expression levels of the linker, core, or variant histones were not statistically different between the two samples (Fig. 3c). Thus, contrary to the previously published study (Berloco *et al*. 2001), we conclude that loss of *ao* does not increase histone gene expression at the steady-state RNA level.

Although not explored in the original study (Berloco *et al*. 2001), we considered the possibility that *ao* might affect histone levels post-transcriptionally. Previous analyses found that histone H2B protein levels are approximately twofold higher in embryos from *ao^1^/ao^1^*females than in embryos from wild-type females, whereas H3 levels are unchanged (Chari *et al*. 2019). To investigate this possibility, we quantified the levels of H2B and H3 proteins in Δ*ao* and isogenic wild-type ovaries using western blotting analyses. We found no evidence for increased histone protein levels in Δ*ao* ovaries; instead, we observed a slight *decrease* in H3 protein levels (Fig. 3d, Table S10).

Thus, our RT-qPCR and western blotting analyses show that loss of *ao* does not significantly increase histone transcript or protein levels in ovaries, challenging previous conclusions that Ao acts as a repressor of core histone expression (Berloco *et al*. 2001).

### Genetic interactions between *ao*, histone genes, and Y-chromosomal heterochromatin

The previous finding that *ao* might encode a histone gene repressor motivated the hypothesis that *ao-*associated maternal-effect lethality results from excess histones in eggs (Berloco *et al*. 2001). This hypothesis led to the prediction that *ao-*associated maternal-effect lethality might be rescued by a histone deficiency in *ao-*mutant mothers. Indeed, a heterozygous deletion of the histone locus in *ao^1^*/*ao^1^* mutant females could ameliorate their maternal-effect lethality (Berloco *et al*. 2001). Although we found no evidence that *ao* encodes a histone gene repressor, we investigated whether a reduction in histone gene copy number could nevertheless suppress the maternal-effect lethality of Δ*ao* females. To test this, we crossed Δ*ao* flies into a strain carrying a heterozygous deletion of the histone locus (BDSC 8670) (Cook *et al*. 2012) to create Δ*ao/*Δ*ao* flies with half the number of histone genes (Fig. S16). We found that reducing histone gene copy number partially rescues the maternal-effect lethality of Δ*ao* females (Fig. 4a), just as previously reported for *ao^1^*/*ao^1^* females (Berloco *et al*. 2001).

**Figure 4.**
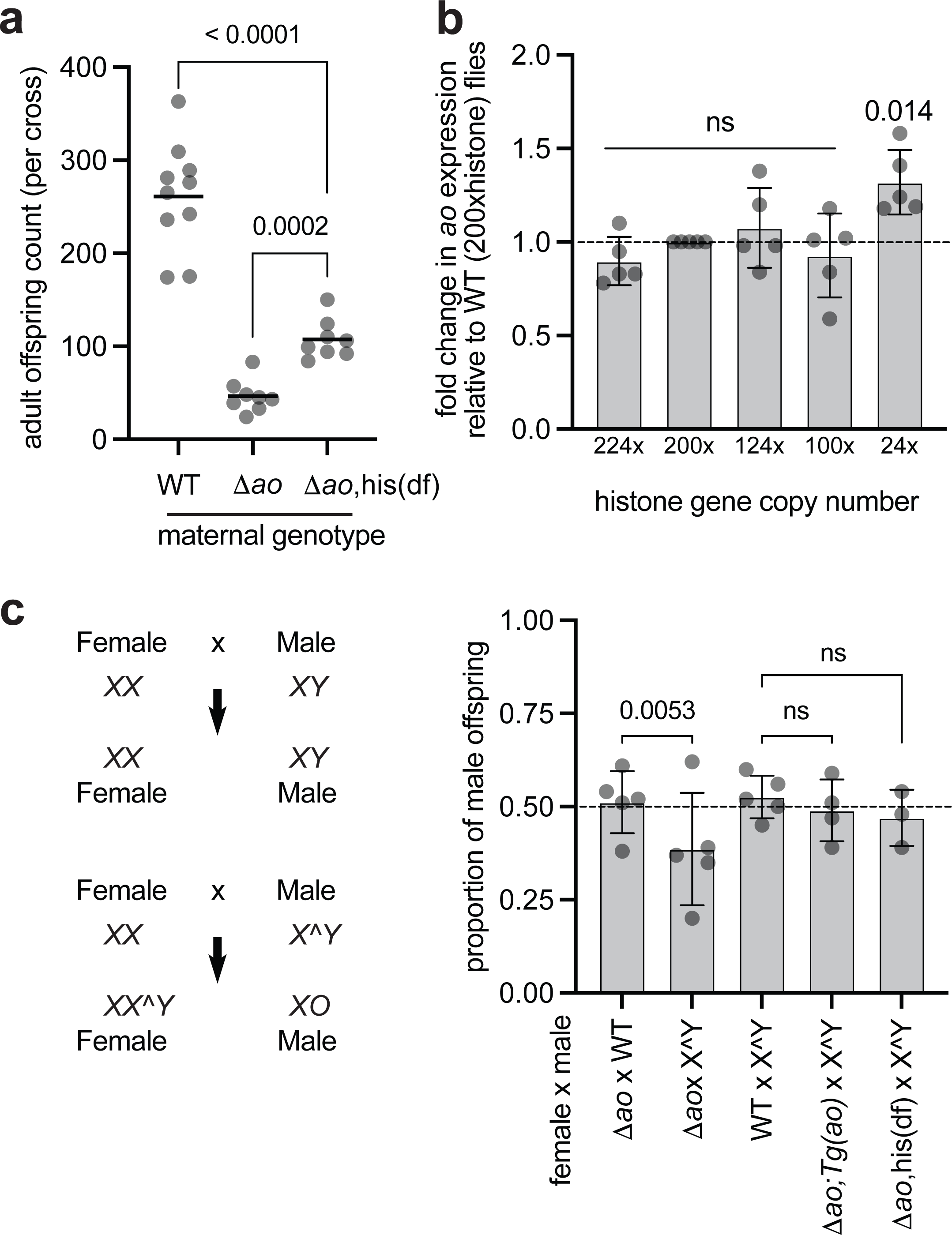
Histone deficiency and Y-chromosome heterochromatin ameliorates *ao’*s maternal-effect lethality. **(a)** We compared the total number of offspring produced by wildtype (isogenic *yw*) females, Δ*ao* females, and Δ*ao* females harboring a hemizygous histone deficiency (*his(df),* which has a deletion from cytological locus 39D3 to 39E2, corresponding to the histone gene array; see Fig. S16) crossed to Δ*ao* males at 29°C. Each point on the graph is the offspring count from a biological replicate cross. The *p-*values are from two-tailed Mann-Whitney U tests. **(b)** Using RT-qPCR, we measured *ao* transcript levels in flies carrying different numbers of histone genes. Each data point is a biological replicate of 4 virgin ovaries. The mean of the biological replicate is shown for each replicate. Gene expression has been normalized to *rp49* and to the wild-type (200xhistone) case. The *p*-values are calculated using a one-sample t-test (Methods). The bars represent one standard deviation. **(c)** Schematic of offspring produced in crosses between XX females and XY males, versus XX females and attached X^Y males. Adult offspring are produced in an equal sex ratio (50% male) in crosses between Δ*ao* females and wildtype males or crosses between wildtype females and attached X^Y males (BDSC strain 9460) at 25°C. In contrast, in crosses between Δ*ao* females and males carrying an attached X^Y sex chromosome, adult progeny counts are skewed to produce more XX^Y females relative to XO males at 25°C. However, this sex-ratio skew is rescued either by the presence of two alleles of the *ao* ‘rescue transgene’ on the 3^rd^ chromosome (Fig. 1c) or by a heterozygous histone deficiency on the 2^nd^ chromosome (Fig. S16). The *p-*values are from a 2×2 contingency table using the two-tailed Fisher’s exact test. The bars represent one standard deviation.

We also assessed the relationship between histone gene copy number and *ao* expression using flies with a deletion of the histone gene cluster (Gunesdogan *et al*. 2010) as well as a ‘12xhistone’ transgene carrying only twelve copies of the histone genes inserted on the 3^rd^ chromosome (Fig. 4b) (Mckay *et al*. 2015). Although wild-type diploid flies encode 200 copies of histone genes, only twelve copies (*i.e.,* flies encoding only a single 12xhistone transgene) can suffice for viability (Gunesdogan *et al*. 2010; Mckay *et al*. 2015). If *ao* were functioning as a histone repressor, we hypothesized that flies with such drastically reduced histone copy number could compensate by reducing *ao* expression, thereby facilitating higher histone gene expression. We used endogenous histone locus deletions and 12xhistone transgenes to generate flies carrying 224, 200 (wild-type), 124, 100, or 24 copies of the histone gene cluster (Fig. S17). Upon quantifying *ao* transcript levels via RT-qPCR, we found that flies encoding 24 copies of histone genes had a nearly 20% increase in *ao* transcript levels compared to flies encoding 100, 124, 200 (wild-type), or 224 copies of the histone genes (Fig. 4b, Table S11). Our findings are consistent with those of a previous study that compared the transcriptomes of 200xhistone and 12xhistone flies (Mcpherson *et al*. 2023). Although it was not the focus of the previous study, this dataset also revealed that *ao* transcript levels were slightly higher in animals with fewer copies of the histone genes. Together, these findings run counter to the expectation that *ao* negatively regulates histone gene expression.

Next, we focused on genetic interactions between *ao* and heterochromatin encoded on sex chromosomes. In crosses involving X^Y fathers, progeny either inherit the paternal attached X^Y chromosome and develop as XX^Y females, or no paternal sex chromosome and develop as (sterile) XO males (Fig. 4c). Although *ao* mutant XX females produce fewer offspring than wild-type females in crosses with XY males due to their maternal-effect lethality, the sex ratio of their offspring is not skewed. However, *ao* mutant females produce an excess number of XX^Y female offspring relative to X0 male offspring when crossed to males with attached X^Y chromosomes (Sandler *et al*. 1968; Sandler 1970; Parry and Sandler 1974; Sandler 1977). This distortion of the offspring sex ratio is attributed to the ability of specific regions of the X- and Y-heterochromatin to partially relieve *ao*-associated maternal-effect lethality (Sandler 1970; Parry and Sandler 1974; Sandler 1977; Yedvobnick *et al*. 1980; Pimpinelli *et al*. 1985; Tomkiel *et al*. 1991). Increased survival of female offspring with Δ*ao* mothers crossed to X^Y fathers would also indicate that the X^Y provides a partial rescue of the Δ*ao* phenotype.

We confirmed the interaction between the *ao* gene and AO heterochromatin on the Y chromosome using the Δ*ao* strain. Crosses between Δ*ao* females and wildtype males yielded nearly equal numbers of female (51%) and male (49%) offspring (Fig. 4c). In contrast, crosses between Δ*ao* females and attached X^Y males (BDSC 9460) yielded a 61:39 offspring ratio of females: males, significantly deviating from the 50:50 expectation (2×2 contingency table, two-tailed Fisher’s exact test, *p*-value=0.0053). Normal sex ratios were restored when Δ*ao* females carrying two copies of the 3^rd^ chromosome ‘rescue’ *ao* transgene were crossed to attached X^Y males (49% males). Similarly, crosses between Δ*ao* females carrying a histone deficiency and attached X^Y males also yielded a nearly equal sex ratio among their progeny (47% males) (Fig. 4c).

Thus, despite our finding that *ao* is not a transcriptional repressor of core histone transcription, our study confirmed previous findings that *ao* loss results in maternal-effect lethality, which can be rescued either by depleting histone gene copy number or by Y chromosome-linked heterochromatin.

## Discussion

Pioneering genetic studies identified *ao* as a maternal-effect lethal gene whose phenotype depended on heterochromatin content in zygotes (Sandler *et al*. 1968; Sandler 1970; Parry and Sandler 1974; Sandler 1977). A satisfying explanation for *ao*’s connection to heterochromatin emerged from a molecular study three decades after its initial characterization, which showed that the *ao*-encoded protein localized to the histone gene cluster and that its loss led to histone overexpression in unfertilized eggs from *ao^1^/ao^2^* mothers (Berloco *et al*. 2001). The same study also demonstrated that reducing histone copy number ameliorated *ao*-associated maternal-effect lethality. These observations led to the model that histone overproduction in *ao/ao* mothers results in high histone levels in eggs and zygotes, which in turn leads to the maternal-effect lethal phenotype. As a result of these findings, *ao* showed promise as a potential tool for manipulating histone gene expression in *Drosophila* (Chari *et al*. 2019), a manipulation that cannot be achieved by simply changing histone gene copy number (Mckay *et al*. 2015; Zhang *et al*. 2019).

Previous studies on *ao* function relied on reagents that have associated genetic background effects. Moreover, two critical reagents – the *ao^2^* allele (Tomkiel *et al*. 1995) and the anti-Ao antibody (Berloco *et al*. 2001) – are no longer available. We developed two novel tools to bridge this gap: a precise CRISPR/Cas9-mediated deletion of the *ao* locus (and replacement with *dsRed*) and *ao* alleles tagged with the V5 epitope at the endogenous locus. Using these reagents, we revisited key findings from the original *ao* studies. We confirmed *ao*-associated maternal-effect lethality and its amelioration via reduced histone copy number or excess Y-heterochromatin.

However, we were unable to replicate two important findings from the previous study (Berloco *et al*. 2001). First, we cannot replicate the previous finding that the Ao protein localizes to the histone gene cluster. This finding could be the result of differences between the specifically-raised, since-lost polyclonal anti-Ao antibody (Berloco *et al*. 2001) and the epitope tags we have used. Nevertheless, we were unable to specifically localize epitope-tagged Ao protein to histone loci in two different tissues (and with two distinct epitope tags in one of the tissues), despite confirming *ao* expression in both tissues. Moreover, we observed no maternal-effect lethality in flies expressing epitope-tagged *ao*-V5, indicating that the epitope tag does not interfere with *ao* function. Second, contrary to the previous study (Berloco *et al*. 2001), we found that *ao* is not a direct repressor of histone expression. For most histone genes, we observed no differences in histone mRNA levels between Δ*ao/*Δ*ao*, *ao^1^/ao^1^*, or wild-type ovaries using multiple assays. Our findings challenge the previously proposed model that histone overexpression results in *ao*’s maternal-effect lethality (Berloco *et al*. 2001).

Since the *ao* study was published, subsequent studies have demonstrated that dramatically reducing histone gene copy number does not decrease histone expression (Mckay *et al*. 2015; Mcpherson *et al*. 2023). These findings also challenge the previous model, which posited that decreased histone gene expression must have compensated for histone overexpression in the absence of *ao*. Since Δ*ao* alleles fully recapitulate the maternal-effect lethality reported in *ao^1^/ao^1^*and *ao^1^/ao^2^*females but not the histone overexpression reported in *ao^1^/ao^2^* trans-heterozygote females, we conclude that histone overexpression is not the mechanism underlying *ao*-associated maternal-effect lethality. Our results further suggest that *ao* should not be used to manipulate histone levels. An alternative tool might be the recently-described histone chaperone Nuclear Autoantigenic Sperm Protein (NASP), which encodes an H3-H4 chaperone in *Drosophila* embryos (Tirgar *et al*. 2023) that prevents H3 aggregation and degradation (Das *et al*. 2025).

Why is there a significant discrepancy in the histone overexpression phenotype between our Δ*ao* alleles and the previously characterized *ao^1^/ao^2^* strain? Precise deletion null alleles (like Δ*ao*) can often differ in phenotype from presumed null alleles that leave part of the gene intact (like *ao^1^*, which has a Doc transposon interrupting the first exon). This difference can result from transcriptional adaptation (El-Brolosy and Stainier 2017), a phenomenon in which alleles that ablate mRNA transcription entirely can be compensated for and, therefore, exhibit less severe phenotypes than those that allow the transcription of mutant mRNA (El-Brolosy *et al*. 2019).

In this case, however, we suspect the explanation here might stem from differences in genetic background among *ao* strains. The recent discovery that histone copy number fluctuates up to five-fold among *Drosophila* strains (Shukla *et al*. 2024) could partially explain the different effects of *ao* alleles on histone levels in different genetic backgrounds observed in previous studies (Berloco *et al*. 2001; Chari *et al*. 2019) and this study. Like in Δ*ao*/Δ*ao* ovaries, we did not observe a histone overexpression phenotype in *ao^1^/ao^1^* ovaries. Before our present study, *ao^1^/ao^1^* flies had never been assessed for histone-transcript levels. Indeed, the overexpression of histone mRNAs had been previously reported only for the *ao^1^/ao^2^* trans-heterozygote (Berloco *et al*. 2001). Unlike the Δ*ao* and *ao^1^* alleles, the *ao^2^* allele could not be made homozygous (Tomkiel *et al*. 1995), indicating the presence of one or more recessive lethal mutations. Moreover, whereas a wild-type *ao* transgene rescues the maternal-effect lethality of *ao^1^* (Tomkiel *et al*. 1995) and Δ*ao* (Fig. 1), a wild-type transgene rescue for histone overexpression or lethality of the *ao^1^*/*ao^2^* trans-heterozygote was never reported (Berloco *et al*. 2001). These observations suggest that the genetic background of the *ao^2^*allele might have conferred an additional phenotypic burden unrelated to *ao* function. For example, *P-*element insertion in the 5’ UTR of *ao* in the *ao^2^* strain might have inadvertently affected the expression of the upstream *ATPsynG* gene, which encodes an essential subunit of the mitochondrial ATP synthase (Fukuoh *et al*. 2014). Unfortunately, since the *ao^2^* strain is no longer available, we cannot test our hypothesis of additional background effects or how they might relate to histone overexpression.

Furthermore, Ao belongs to the DET1 family of E3 ubiquitin ligases (Berloco *et al*. 2001). Orthologs of *ao/DET1* are present as single-copy genes in most plant and animal genomes, including human *hDET1*, a negative regulator of a proto-oncogene (Wertz *et al*. 2004; Pick *et al*. 2007), and *Arabidopsis DET1,* which functions as a negative regulator of light-mediated growth in seedlings (Chory *et al*. 1989; Chory and Peto 1990; Pepper *et al*. 1994). *Arabidopsis* DET1 binds to nonacetylated, C-terminal H2B tails in the nucleosome (Benvenuto *et al*. 2002) and regulates H2B mono-ubiquitination in a light-dependent context (Nassrallah *et al*. 2018). Given the conservation of the plant and mammalian DET1 proteins as subunits of the COP1 Cul4A-RING E3 ubiquitin ligase complex (Wertz *et al*. 2004; Yanagawa *et al*. 2004; Bernhardt *et al*. 2006; Pick *et al*. 2007), it is highly likely that *Drosophila* Ao also functions in post-translational, rather than transcriptional, regulation. However, our western blotting analyses suggest that histones are not the post-translational target of Ao.

Our findings suggest that a molecular mechanism distinct from histone gene repression underlies *ao* function, its maternal-effect lethality, and its genetic interactions with the histone gene cluster and heterochromatin. Based on its homology and genetic interactions, we hypothesize that Ao functions as a DET1-family E3 ligase adaptor that post-translationally regulates a dosage-sensitive chromatin-associated factor, whose dosage sensitivity is buffered by both the histone locus and sex-chromosome heterochromatin. Under this model, Ao might target a component of the histone locus body (HLB), which regulates histone biogenesis (White *et al*. 2011; Duronio and Marzluff 2017), or a tightly associated chromatin factor, for ubiquitin-dependent degradation. In wild-type flies, Ao keeps this factor at an optimal level, allowing normal HLB assembly and histone mRNA biogenesis, and permitting embryonic development across a wide range of histone copy numbers and heterochromatin contents. However, loss of Ao would cause this factor to become overabundant or misregulated at ‘vulnerable’ repetitive regions, disrupting early embryonic nuclear functions (*e.g.*, replication timing, chromatin packaging, chromosome segregation) and resulting in maternal-effect lethality. In this model, loss of *ao* could be partially compensated either by 1) simultaneous depletion of this unknown component via reduction or loss of the histone locus, leading to its destabilization independent of Ao, or 2) sequestration of excess levels of this factor at sex-chromosome heterochromatin sinks. An alternative possibility is that the histone gene cluster might serve as a sink for another factor that becomes essential for embryonic viability in the absence of *ao*. When histone gene dosage is reduced, this factor is freed to relieve maternal-effect lethality caused by loss of *ao*. Under this model, *D. melanogaster* strains that naturally encode high histone gene copy number (Shukla *et al*. 2024) might be prone to more severe maternal-effect lethality upon loss of *ao*, whereas strains that naturally encode fewer genes or those carrying a histone deficiency might be better able to withstand the loss of *ao*. Although our findings challenge the current molecular model of *ao*, these possibilities highlight that understanding the basis of *ao*-associated maternal-effect lethality and its connection to histone copy number and heterochromatin remains an open and exciting question.

## Acknowledgments

We thank Sarah Tomlin for assistance with the paternal-effect fertility assays, Kaitlin Koreski for help with Sanger sequencing to confirm the transgenic flies, and L. Aravind for discussions on DET1 homology. We further thank Ching-Ho Chang, Akhila Rajan, and Courtney Schroeder for their experimental advice, and the members of the Rieder and Malik labs for their valuable discussions. We would like to extend our special thanks to Richard McLaughlin and Mosur Raghuraman for their valuable comments on the manuscript. Various plasmids used in the study were obtained from the *Drosophila* Genomics Resource Center (NIH grant 2P40OD010949) and Addgene. We thank Flybase (NIH grant U24 HG013300) for maintaining up-to-date information on *D. melanogaster* genes, and Robert Duronio for the kind gift of the Mxc antibody and the V5-Lsm11 strain. We dedicate our study to Larry Sandler and Sergio Pimpinelli for their pioneering work on *Drosophila* heterochromatin.

## Data availability statement

All data for the paper are available in the supplementary materials. RNA-seq data is available in NCBI’s Sequence Read Archive (SRA) under the BioProject ID of PRJNA1199081.

## References

Benvenuto, G., F. Formiggini, P. Laflamme, M. Malakhov and C. Bowler, 2002 The photomorphogenesis regulator DET1 binds the amino-terminal tail of histone H2B in a nucleosome context. Curr Biol 12: 1529–1534.

Berloco, M., L. Fanti, A. Breiling, V. Orlando and S. Pimpinelli, 2001 The maternal effect gene, abnormal oocyte (abo), of Drosophila melanogaster encodes a specific negative regulator of histones. Proceedings of the National Academy of Sciences 98: 12126–12131.

Bernhardt, A., E. Lechner, P. Hano, V. Schade, M. Dieterle et al., 2006 CUL4 associates with DDB1 and DET1 and its downregulation affects diverse aspects of development in Arabidopsis thaliana. Plant J 47: 591–603.

Bischof, J., M. Björklund, E. Furger, C. Schertel, J. Taipale et al., 2013 A versatile platform for creating a comprehensive UAS-ORFeome library in Drosophila. Development 140: 2434–2442.

Brown, J. B., N. Boley, R. Eisman, G. E. May, M. H. Stoiber et al., 2014 Diversity and dynamics of the Drosophila transcriptome. Nature 512: 393–399.

Bulchand, S., S. D. Menon, S. E. George and W. Chia, 2010 Muscle wasted: a novel component of the Drosophila histone locus body required for muscle integrity. J Cell Sci 123: 2697–2707.

Cavaliere, V., F. Graziani, S. Andone, A. Manzi and C. Malva, 1991 Complete reversion of the abo phenotype in D. melanogaster occurs only when the blood transposon is lost from region 32E. Mol Gen Genet 230: 433–441.

Chari, S., H. Wilky, J. Govindan and A. A. Amodeo, 2019 Histone concentration regulates the cell cycle and transcription in early development. Development 146: dev177402.

Chory, J., C. Peto, R. Feinbaum, L. Pratt and F. Ausubel, 1989 Arabidopsis thaliana mutant that develops as a light-grown plant in the absence of light. Cell 58: 991–999.

Chory, J., and C. A. Peto, 1990 Mutations in the DET1 gene affect cell-type-specific expression of light-regulated genes and chloroplast development in Arabidopsis. Proc Natl Acad Sci U S A 87: 8776–8780.

Cook, R. K., S. J. Christensen, J. A. Deal, R. A. Coburn, M. E. Deal et al., 2012 The generation of chromosomal deletions to provide extensive coverage and subdivision of the Drosophila melanogaster genome. Genome biology 13: 1–14.

Das, M., E. Coronado-Chavez, A. D. Bhatt, R. Tirgar, A. A. Amodeo et al., 2025 NASP functions in the cytoplasm to prevent histone H3 aggregation during early embryogenesis. bioRxiv.

Duronio, R. J., and W. F. Marzluff, 2017 Coordinating cell cycle-regulated histone gene expression through assembly and function of the Histone Locus Body. RNA Biol 14: 726–738.

El-Brolosy, M. A., Z. Kontarakis, A. Rossi, C. Kuenne, S. Günther et al., 2019 Genetic compensation triggered by mutant mRNA degradation. Nature 568: 193–197.

El-Brolosy, M. A., and D. Y. Stainier, 2017 Genetic compensation: A phenomenon in search of mechanisms. PLoS genetics 13: e1006780.

Fukuoh, A., G. Cannino, M. Gerards, S. Buckley, S. Kazancioglu et al., 2014 Screen for mitochondrial DNA copy number maintenance genes reveals essential role for ATP synthase. Mol Syst Biol 10: 734.

Godfrey, A. C., A. E. White, D. C. Tatomer, W. F. Marzluff and R. J. Duronio, 2009 The Drosophila U7 snRNP proteins Lsm10 and Lsm11 are required for histone pre-mRNA processing and play an essential role in development. RNA 15: 1661–1672.

Gratz, S. J., F. P. Ukken, C. D. Rubinstein, G. Thiede, L. K. Donohue et al., 2014 Highly specific and efficient CRISPR/Cas9-catalyzed homology-directed repair in Drosophila. Genetics 196: 961–971.

Graziani, F., L. Vicari, E. Boncinelli, C. Malva, A. Manzi et al., 1981 Selective replication of ribosomal DNA repeats after loss of the abnormal oocyte phenotype in Drosophila melanogaster. Proc Natl Acad Sci U S A 78: 7662–7664.

Groth, A. C., M. Fish, R. Nusse and M. P. Calos, 2004 Construction of transgenic Drosophila by using the site-specific integrase from phage phiC31. Genetics 166: 1775–1782.

Gunesdogan, U., H. Jackle and A. Herzig, 2010 A genetic system to assess in vivo the functions of histones and histone modifications in higher eukaryotes. EMBO Rep 11: 772–776.

Haemer, J., 1978 Studies on heterochromatin of Drosophila melanogaster, pp. in Genetics. University of Washington.

Hodkinson, L. J., J. Gross, C. A. Schmidt, P. P. Diaz-Saldana, T. Aoki et al., 2024 Sequence reliance of the Drosophila context-dependent transcription factor CLAMP. Genetics 227.

Kim, D., J. M. Paggi, C. Park, C. Bennett and S. L. Salzberg, 2019 Graph-based genome alignment and genotyping with HISAT2 and HISAT-genotype. Nat Biotechnol 37: 907–915.

Kremer, H., and W. Hennig, 1990 Isolation and characterization of a Drosophila hydei histone DNA repeat unit. Nucleic acids research 18: 1573–1586.

Krider, H. M., and B. I. Levine, 1975 Studies on the mutation abnormal oocyte and its interaction with the ribosomal DNA of Drosophila melanogaster. Genetics 81: 501–513.

Krider, H. M., B. Yedvobnick and B. I. Levine, 1979 The effect of abo phenotypic expression on ribosomal DNA instabilities in Drosophila melanogaster. Genetics 92: 879–889.

Lifton, R., M. Goldberg, R. Karp and D. Hogness, 1978 The organization of the histone genes in Drosophila melanogaster: functional and evolutionary implications, pp. 1047-1051 in Cold Spring Harbor symposia on quantitative biology. Cold Spring Harbor Laboratory Press.

Livak, K. J., and T. D. Schmittgen, 2001 Analysis of relative gene expression data using real-time quantitative PCR and the 2(-Delta Delta C(T)) Method. Methods 25: 402–408.

Love, M. I., W. Huber and S. Anders, 2014 Moderated estimation of fold change and dispersion for RNA-seq data with DESeq2. Genome Biol 15: 550.

Manzi, A., F. Graziani, T. Labella, G. Gargiulo, F. Rafti et al., 1986 Changes in abo phenotypic expression without increase in rDNA in Drosophila melanogaster. Mol Gen Genet 205: 366–371.

Marzluff, W. F., E. J. Wagner and R. J. Duronio, 2008 Metabolism and regulation of canonical histone mRNAs: life without a poly(A) tail. Nat Rev Genet 9: 843–854.

McKay, D. J., S. Klusza, T. J. Penke, M. P. Meers, K. P. Curry et al., 2015 Interrogating the function of metazoan histones using engineered gene clusters. Developmental cell 32: 373–386.

McPherson, J. E., L. C. Grossmann, H. R. Salzler, R. L. Armstrong, E. Kwon et al., 2023 Reduced histone gene copy number disrupts Drosophila Polycomb function. Genetics 224.

Nassrallah, A., M. Rougée, C. Bourbousse, S. Drevensek, S. Fonseca et al., 2018 DET1-mediated degradation of a SAGA-like deubiquitination module controls H2Bub homeostasis. Elife 7: e37892.

Ozturk-Colak, A., S. J. Marygold, G. Antonazzo, H. Attrill, D. Goutte-Gattat et al., 2024 FlyBase: updates to the Drosophila genes and genomes database. Genetics 227.

Parry, D. M., and L. Sandler, 1974 The genetic identification of a heterochromatic segment on the X chromosome of Drosophila melanogaster. Genetics 77: 535–539.

Pepper, A., T. Delaney, T. Washburnt, D. Poole and J. Chory, 1994 DET1, a negative regulator of light-mediated development and gene expression in Arabidopsis, encodes a novel nuclear-localized protein. Cell 78: 109–116.

Pick, E., O.-S. Lau, T. Tsuge, S. Menon, Y. Tong et al., 2007 Mammalian DET1 regulates Cul4A activity and forms stable complexes with E2 ubiquitin-conjugating enzymes. Molecular and cellular biology 27: 4708–4719.

Pimpinelli, S., W. Sullivan, M. Prout and L. Sandler, 1985 On biological functions mapping to the heterochromatin of Drosophila melanogaster. Genetics 109: 701–724.

Port, F., and S. L. Bullock, 2016 Augmenting CRISPR applications in Drosophila with tRNA-flanked sgRNAs. Nature methods 13: 852–854.

Port, F., H.-M. Chen, T. Lee and S. L. Bullock, 2014 Optimized CRISPR/Cas tools for efficient germline and somatic genome engineering in Drosophila. Proceedings of the National Academy of Sciences 111: E2967–E2976.

Quinlan, A. R., and I. M. Hall, 2010 BEDTools: a flexible suite of utilities for comparing genomic features. Bioinformatics 26: 841–842.

Rieder, L. E., K. P. Koreski, K. A. Boltz, G. Kuzu, J. A. Urban et al., 2017 Histone locus regulation by the Drosophila dosage compensation adaptor protein CLAMP. Genes Dev 31: 1494–1508.

Sandler, L., 1970 The regulation of sex chromosome heterochromatic activity by an autosomal gene in Drosophila melanogaster. Genetics 64: 481.

Sandler, L., 1977 Evidence for a set of closely linked autosomal genes that interact with sex-chromosome heterochromatin in Drosophila melanogaster. Genetics 86: 567–582.

Sandler, L., D. Lindsley, B. Nicoletti and G. Trippa, 1968 Mutants affecting meiosis in natural populations of Drosophila melanogaster. Genetics 60: 525.

Shukla, H. G., M. Chakraborty and J. J. Emerson, 2024 Genetic variation in recalcitrant repetitive regions of the Drosophila melanogaster genome. bioRxiv.

Spring, A. M., A. C. Raimer, C. D. Hamilton, M. J. Schillinger and A. G. Matera, 2019 Comprehensive Modeling of Spinal Muscular Atrophy in Drosophila melanogaster. Front Mol Neurosci 12: 113.

Strausbaugh, L. D., and E. S. Weinberg, 1982 Polymorphism and stability in the histone gene cluster of Drosophila melanogaster. Chromosoma 85: 489–505.

Sullivan, W., and S. Pimpinelli, 1986 The genetic factors altered in homozygous abo stocks of Drosophila melanogaster. Genetics 114: 885–895.

Tirgar, R., J. P. Davies, L. Plate and J. T. Nordman, 2023 The histone chaperone NASP maintains H3-H4 reservoirs in the early Drosophila embryo. PLoS Genet 19: e1010682.

Tomkiel, J., L. Fanti, M. Berloco, L. Spinelli, J. W. Tamkun et al., 1995 Developmental genetical analysis and molecular cloning of the abnormal oocyte gene of Drosophila melanogaster. Genetics 140: 615–627.

Tomkiel, J., S. Pimpinelli and L. Sandler, 1991 Rescue from the abnormal oocyte maternal-effect lethality by ABO heterochromatin in Drosophila melanogaster. Genetics 128: 583–594.

Venken, K. J., Y. He, R. A. Hoskins and H. J. Bellen, 2006 P [acman]: a BAC transgenic platform for targeted insertion of large DNA fragments in D. melanogaster. Science 314: 1747–1751.

Wertz, I. E., K. M. O’Rourke, Z. Zhang, D. Dornan, D. Arnott et al., 2004 Human De-etiolated-1 regulates c-Jun by assembling a CUL4A ubiquitin ligase. Science 303: 1371–1374.

White, A. E., B. D. Burch, X. C. Yang, P. Y. Gasdaska, Z. Dominski et al., 2011 Drosophila histone locus bodies form by hierarchical recruitment of components. J Cell Biol 193: 677–694.

Yanagawa, Y., J. A. Sullivan, S. Komatsu, G. Gusmaroli, G. Suzuki et al., 2004 Arabidopsis COP10 forms a complex with DDB1 and DET1 in vivo and enhances the activity of ubiquitin conjugating enzymes. Genes Dev 18: 2172–2181.

Yedvobnick, B., H. M. Krider and B. I. Levine, 1980 Analysis of the autosomal mutation abo and its interaction with the ribosomal DNA or Drosophila melanogaster: the role of X-chromosome heterochromatin. Genetics 95: 661–672.

Zhang, W., X. Zhang, Z. Xue, Y. Li, Q. Ma et al., 2019 Probing the function of metazoan histones with a systematic library of H3 and H4 mutants. Developmental cell 48: 406–419. e405.

