## Supplementary figures and images for "The *Drosophila* maternal-effect gene *abnormal oocyte* (*ao*) does not repress histone gene expression"

### Figure S1

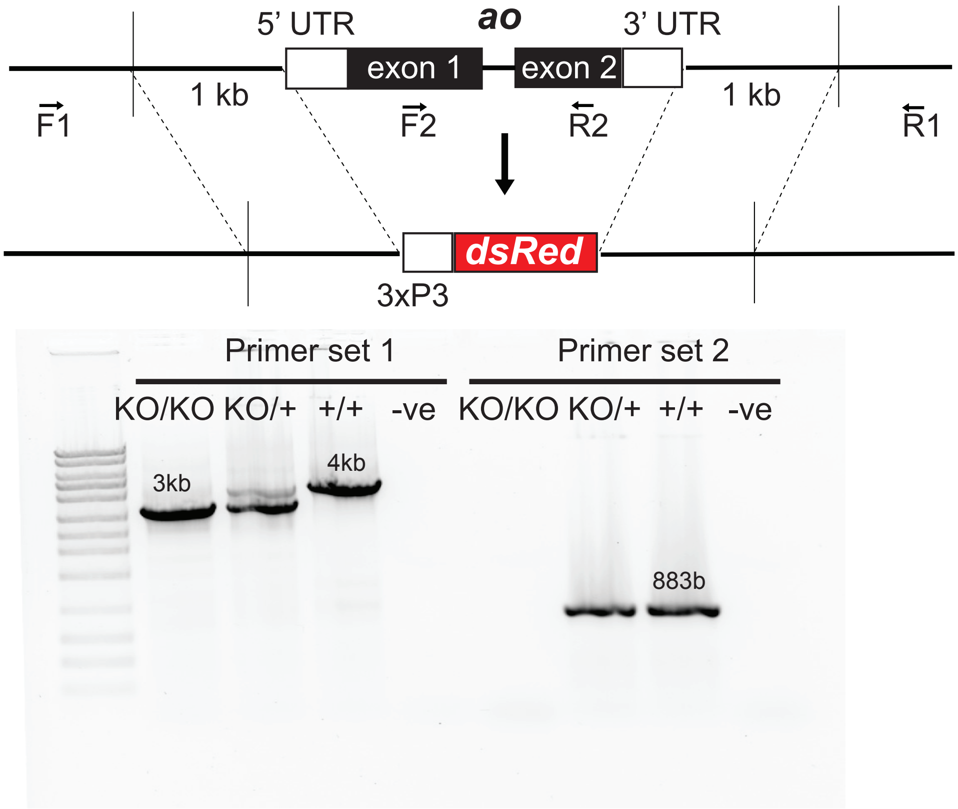

### Figure S3

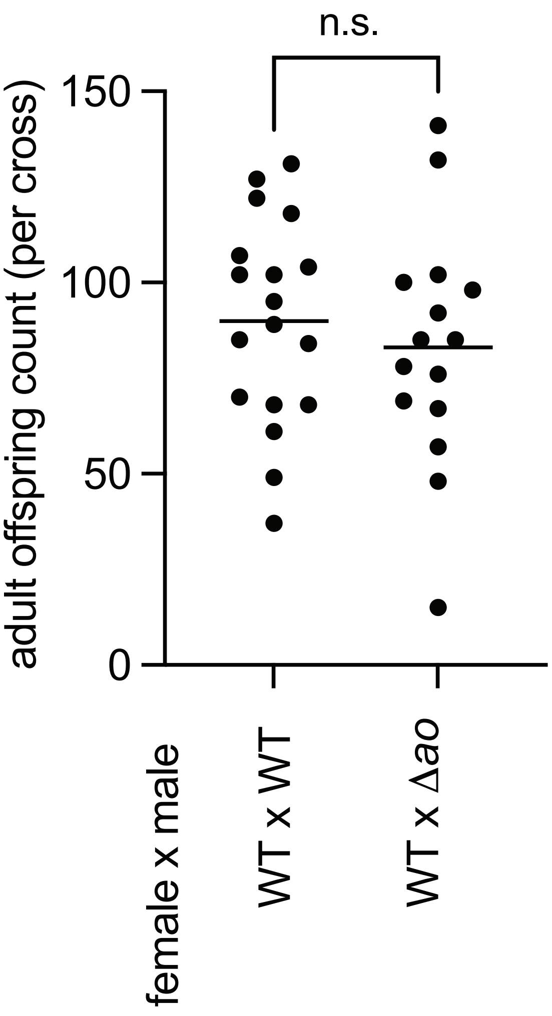

### Figure S4

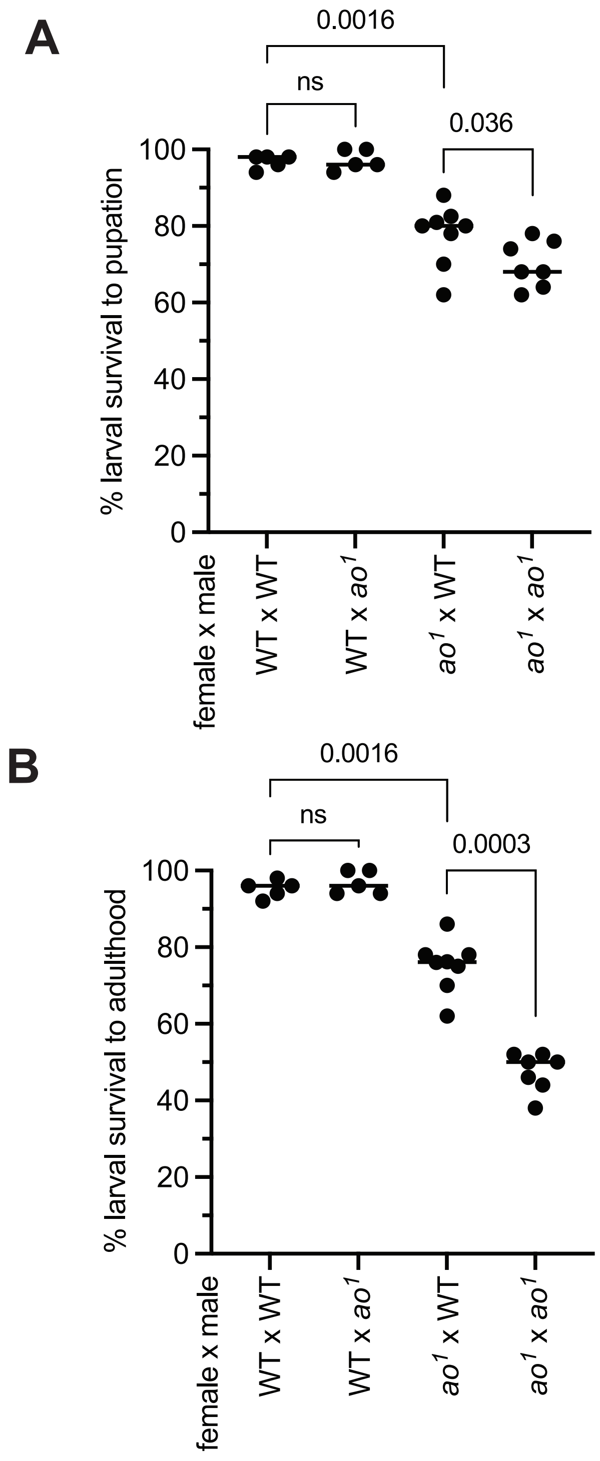

### Figure S5

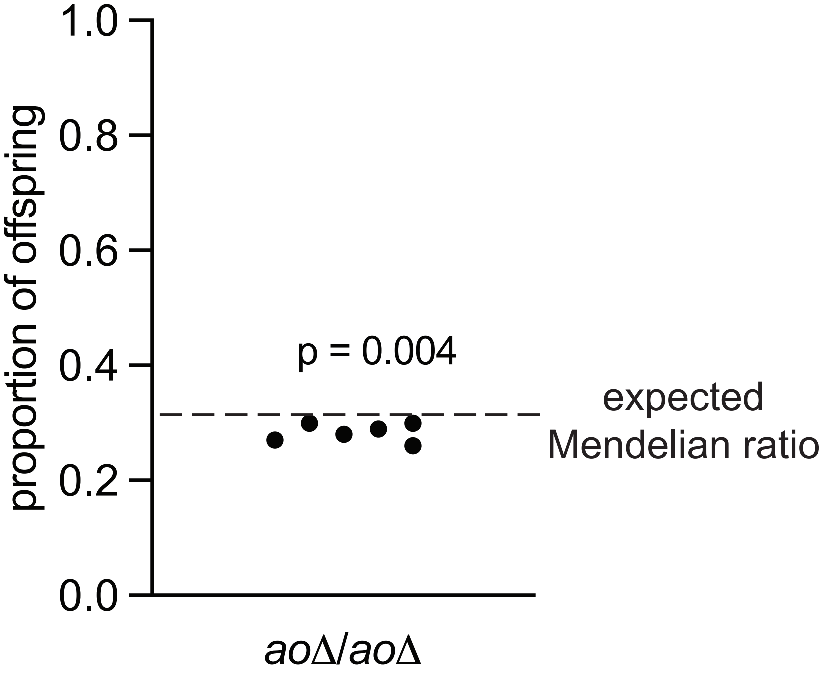

### Figure S6

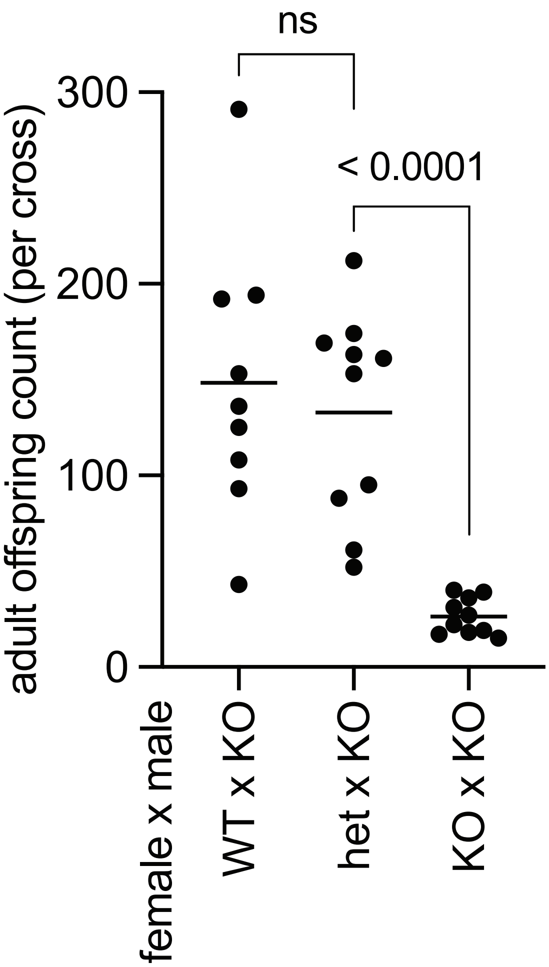

### Figure S7

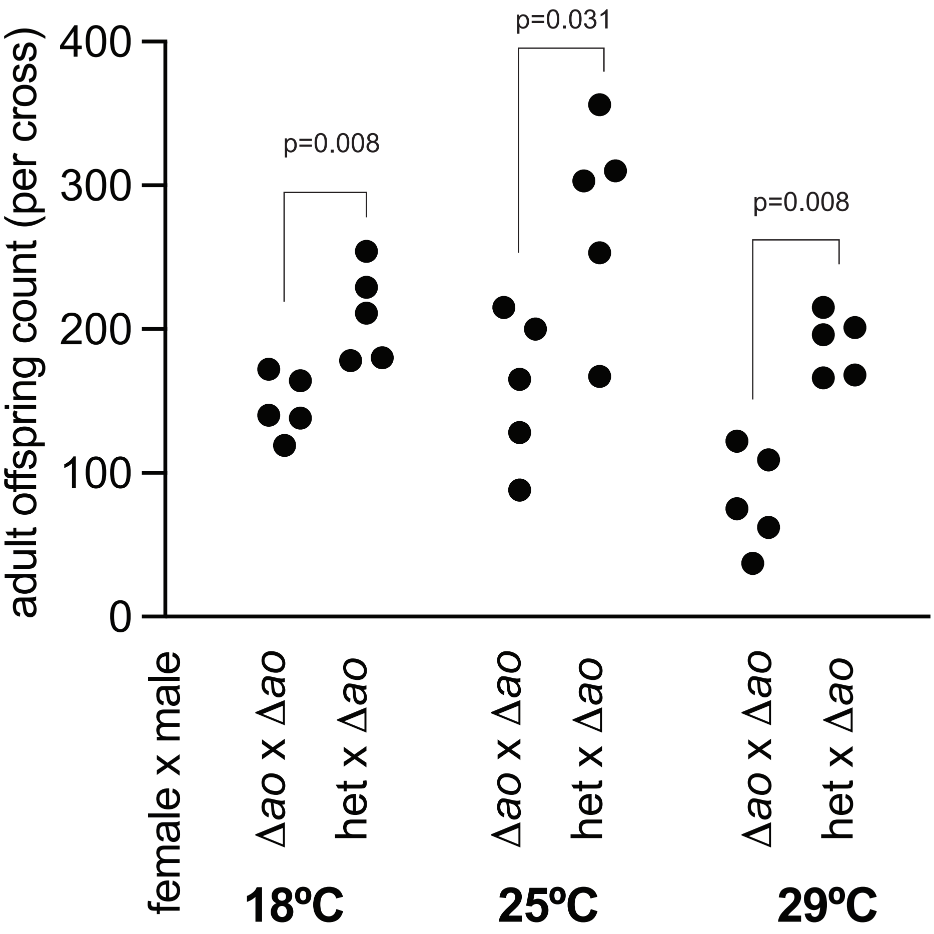

### Figure S8

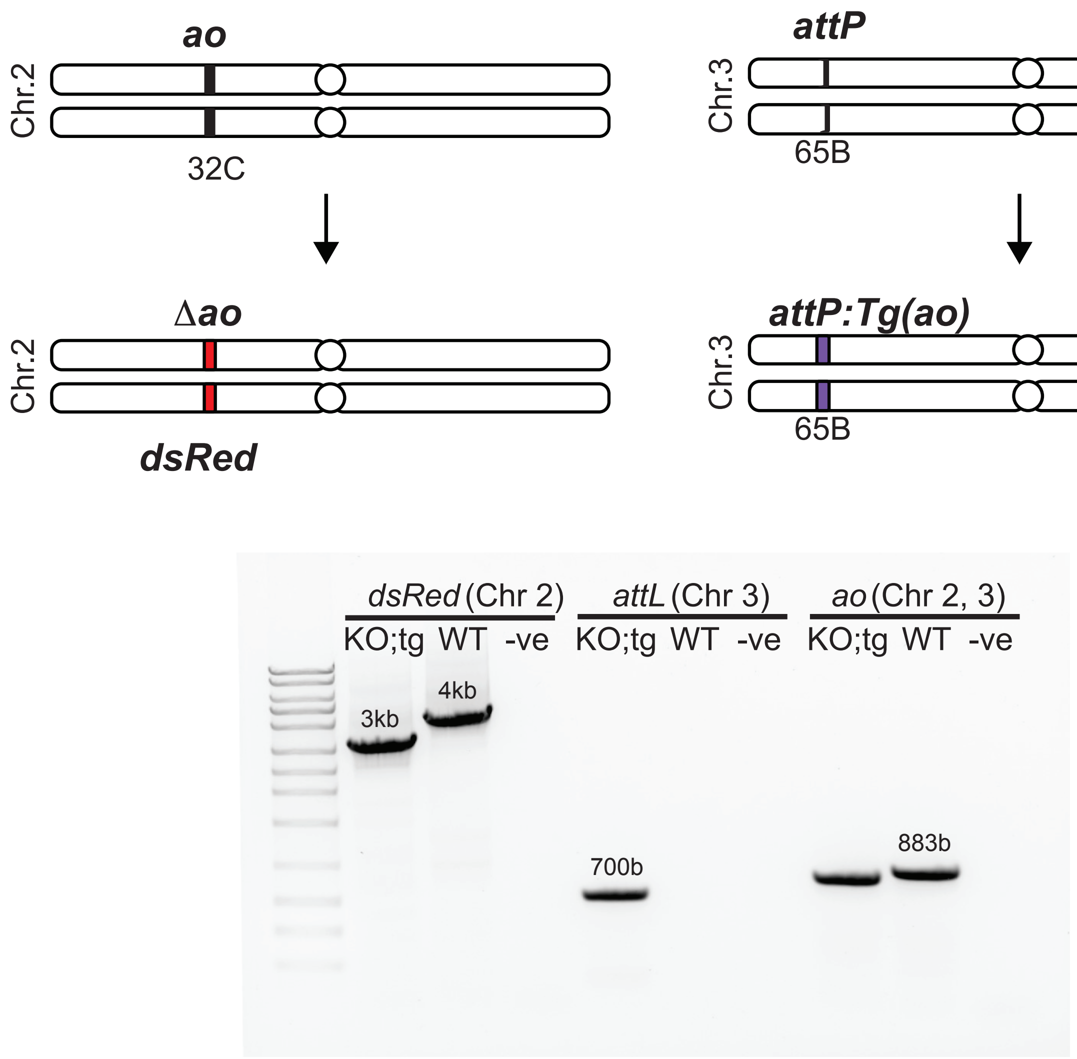

### Figure S9

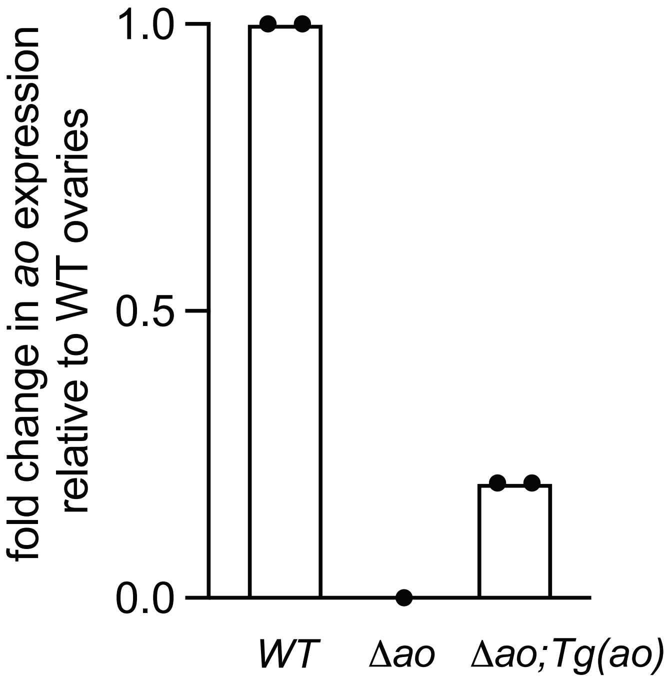

### Figure S10

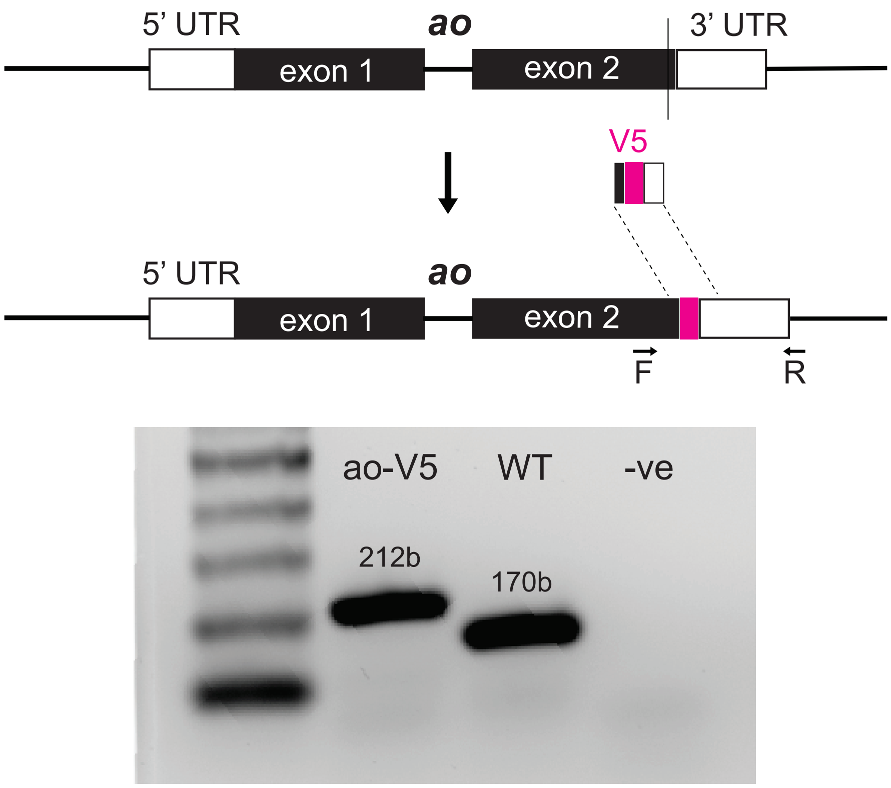

### Figure S11

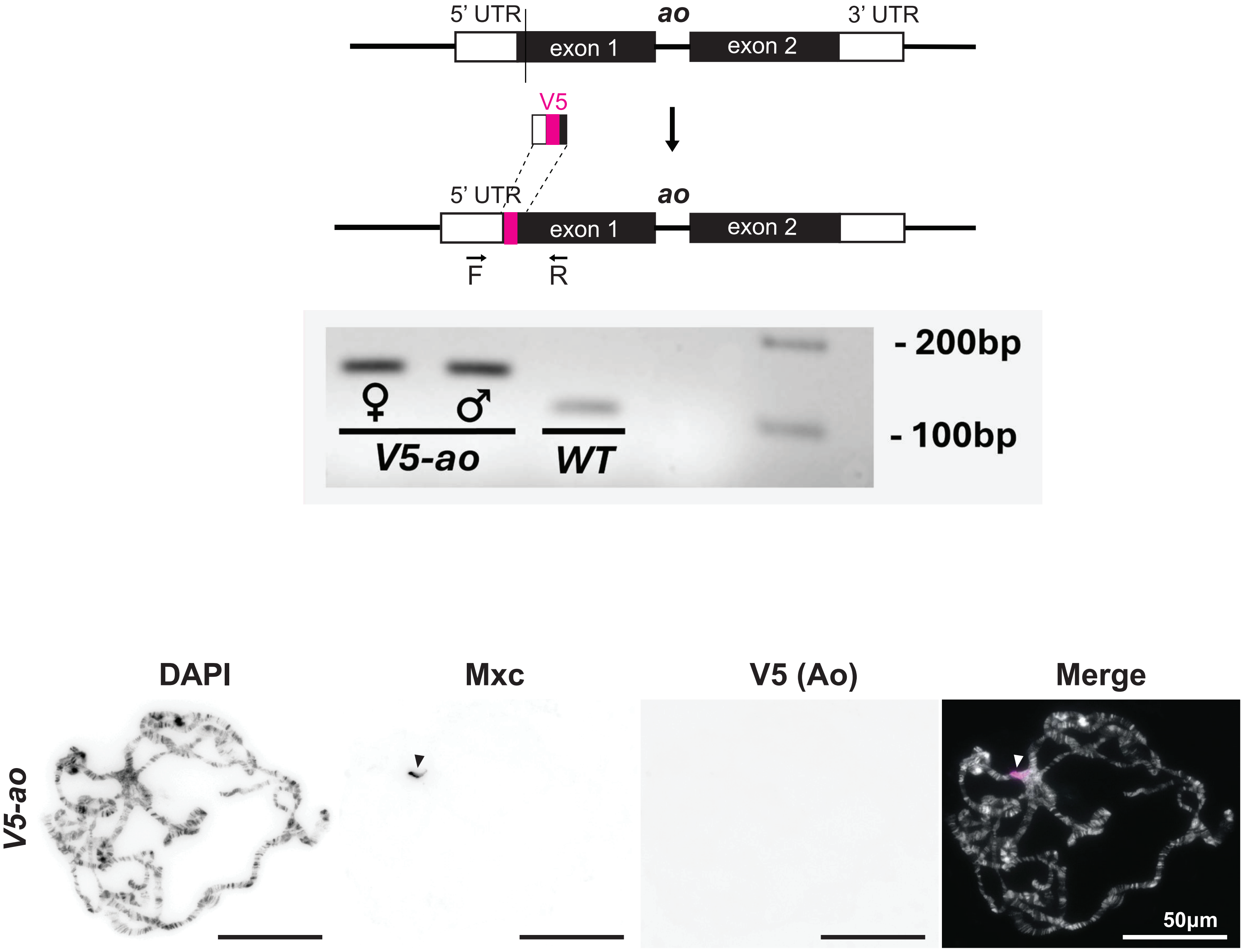

### Figure S12

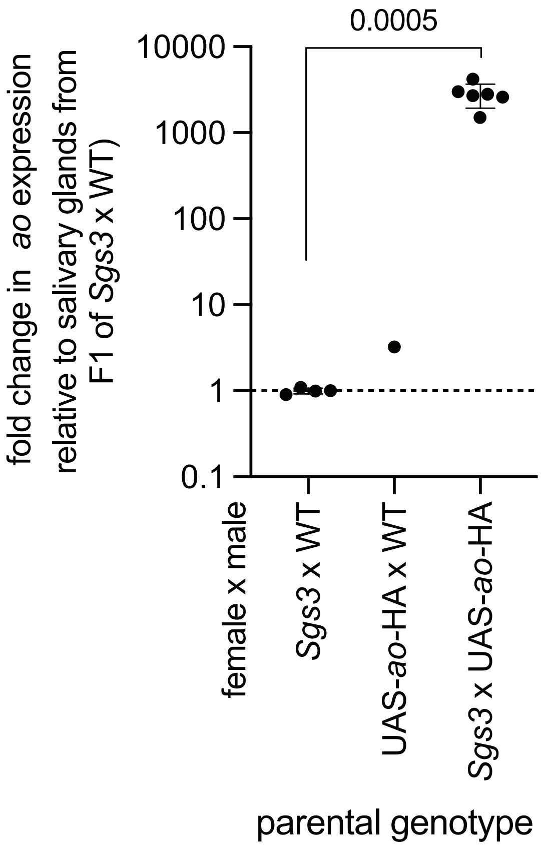

### Figure S13

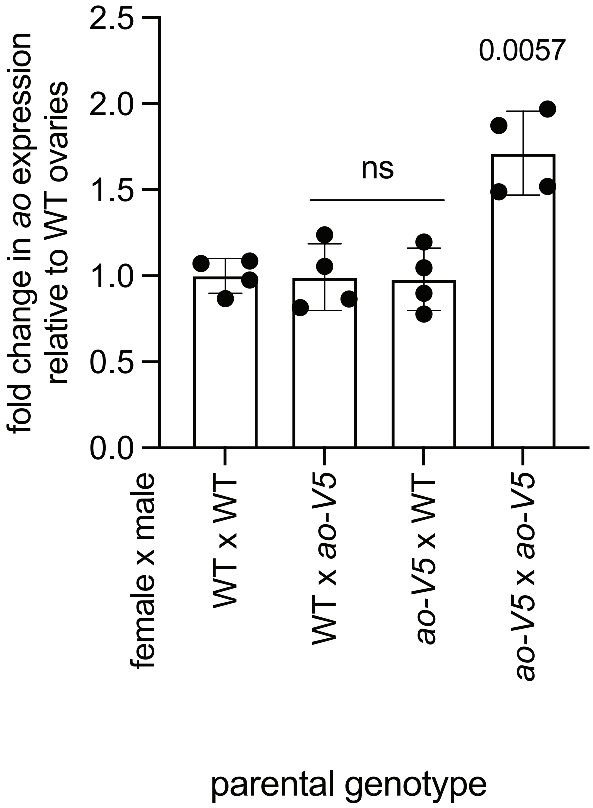

### Figure S14

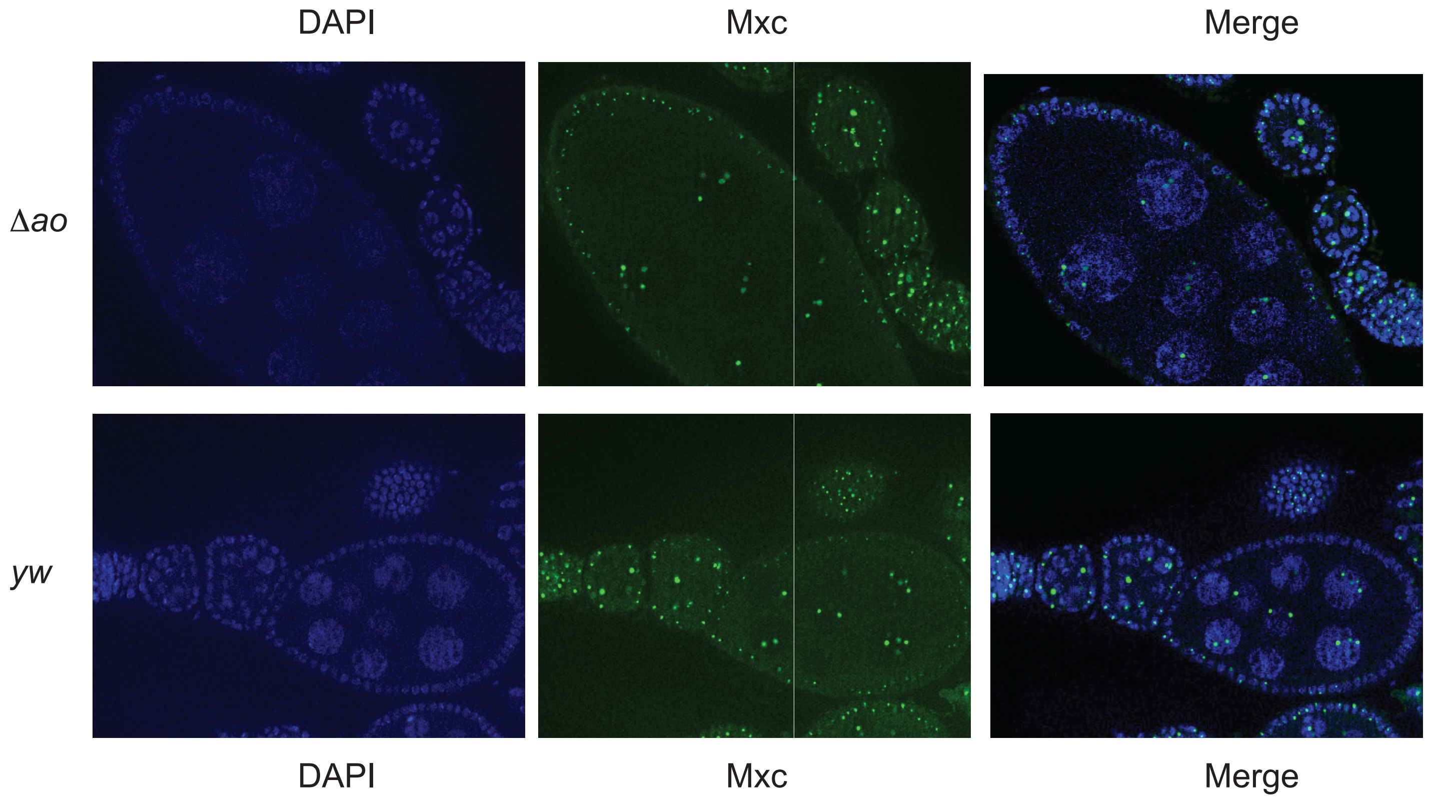


**Figure S14**

### Figure S15

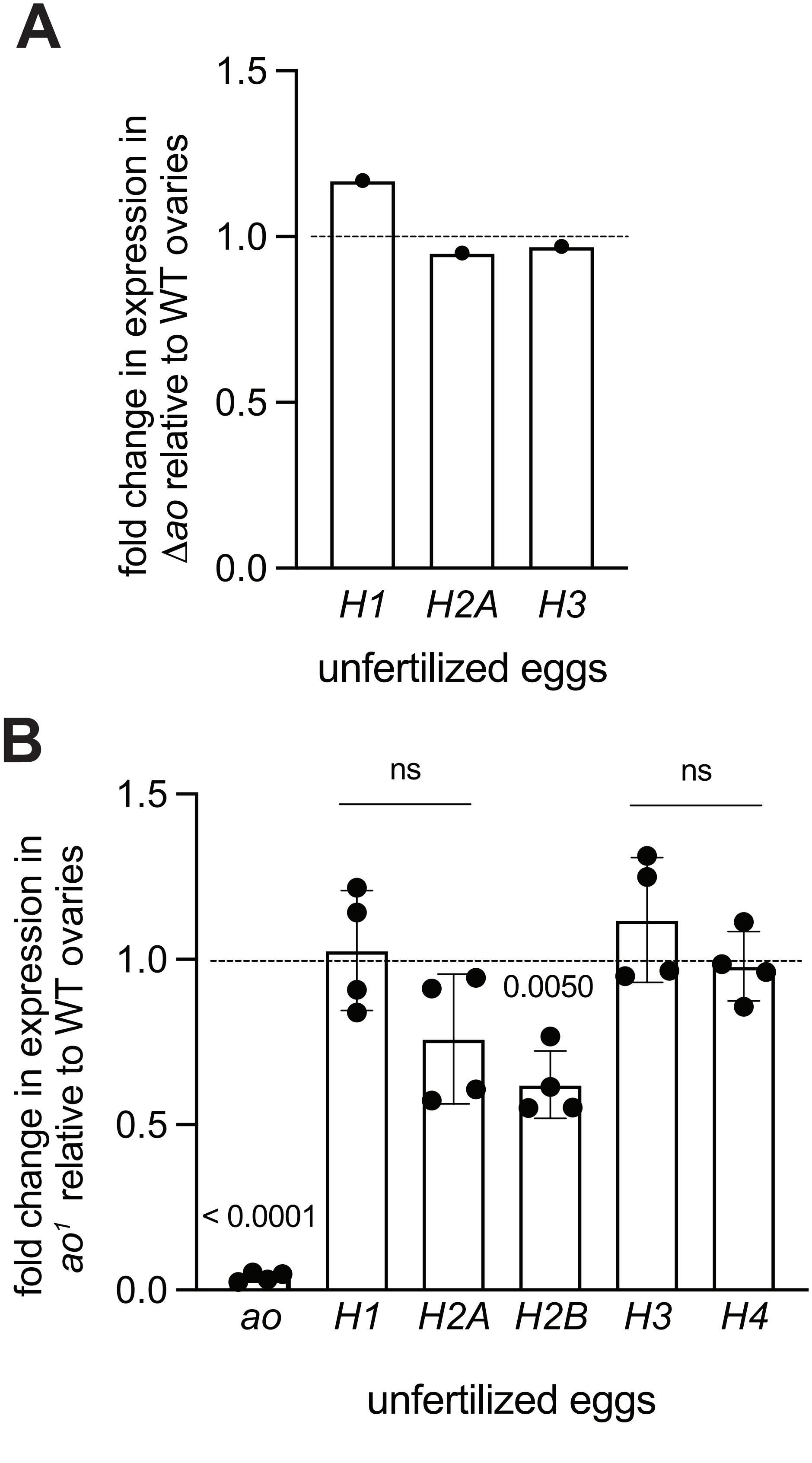

### Figure S16

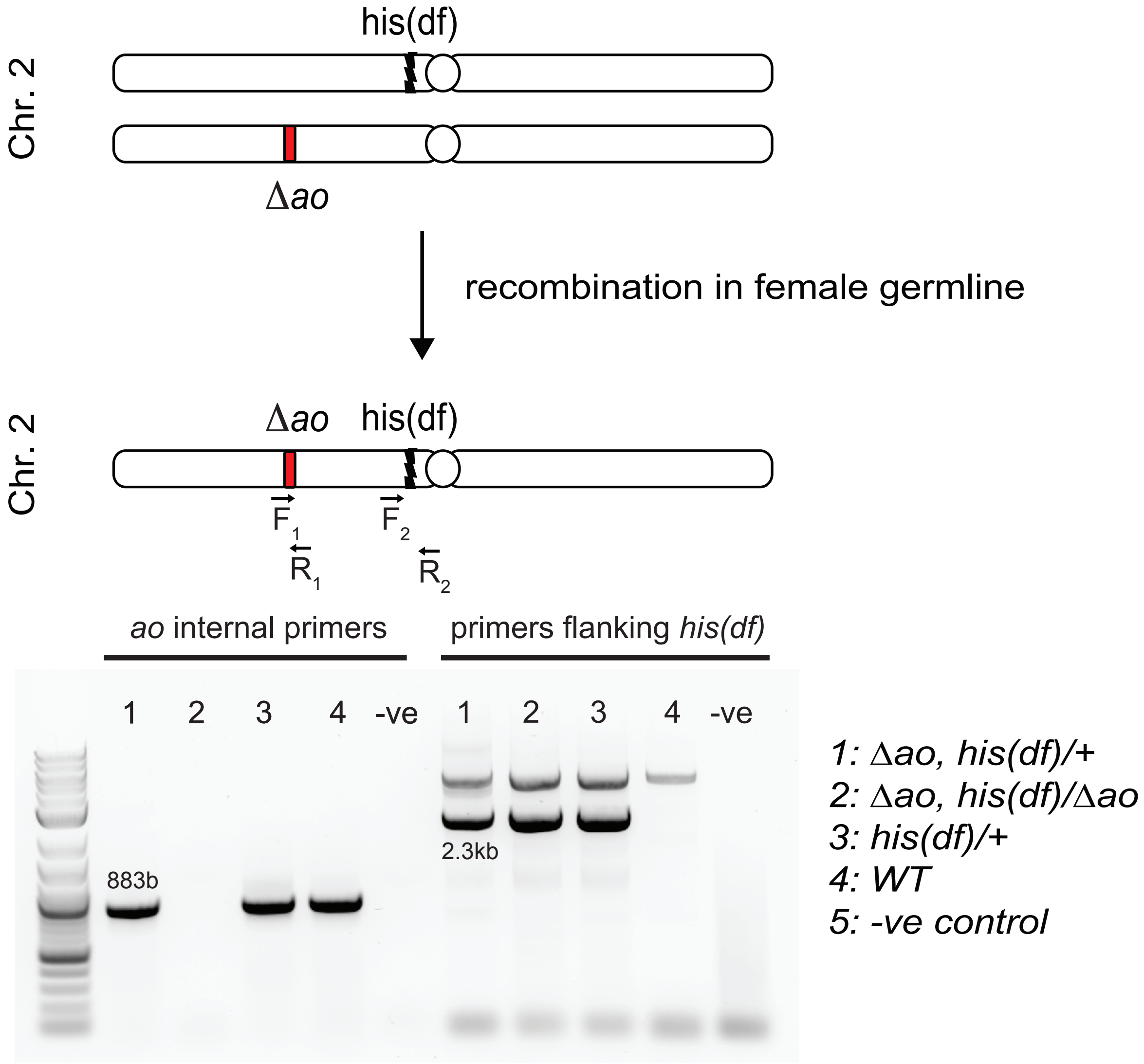

### Figure S17

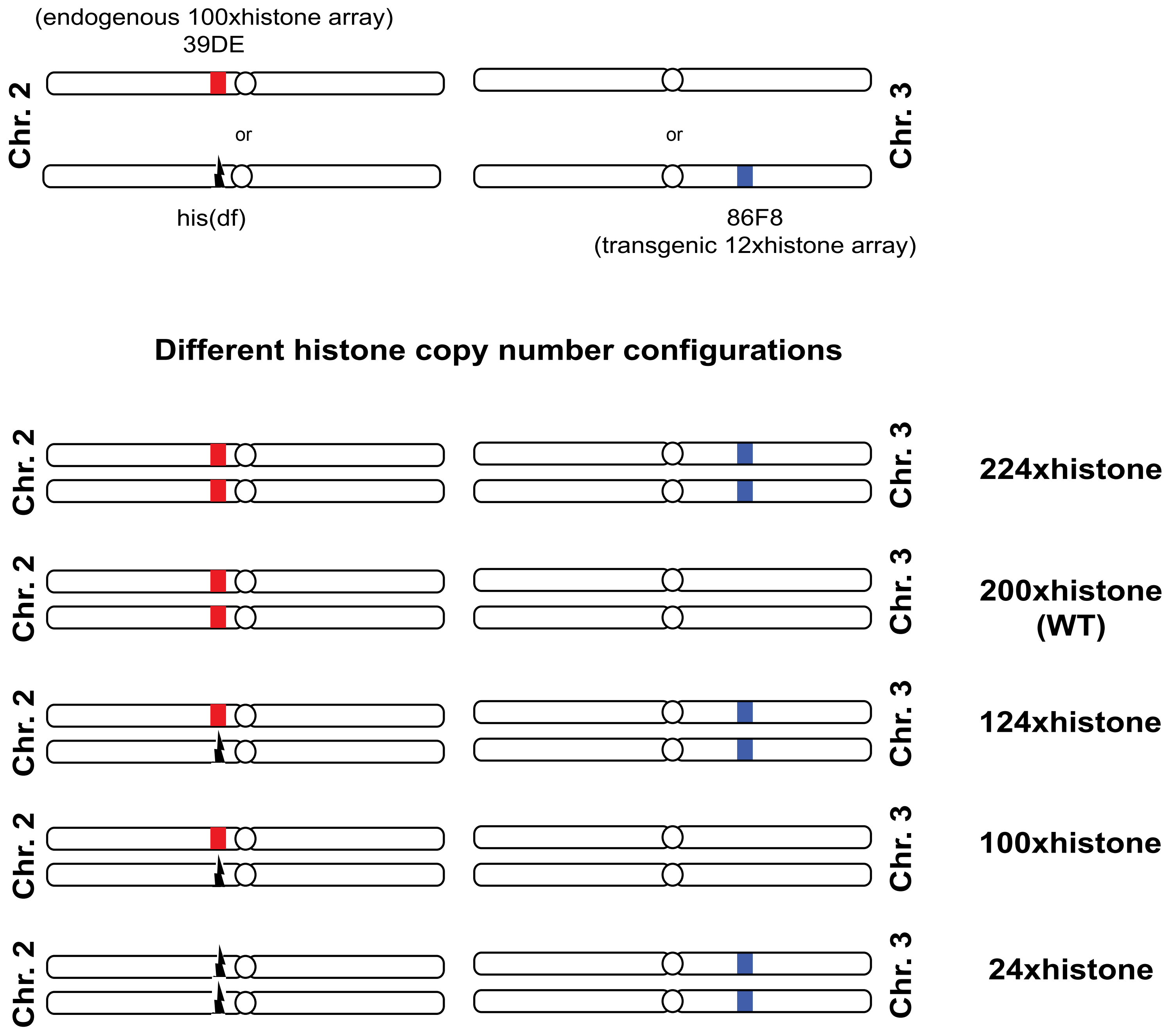
