## Supplementary material for "The *Drosophila* maternal-effect gene *abnormal oocyte* (*ao*) does not repress histone gene expression": Figure S2

( | = identical between Δ*ao* and isogenic *yw* strain)

```
                      ATP synthase subunit G
CAAGGTGGAACTGACGCCCCCGACGCCCGCCGATATTCCGGCCATTCGCCAAGGACTGGG
||||||||||||||||||||||||||||||||||||||||||||||||||||||||||||

CAACATCATCAAGGGAGCCAAGACCGGCGCCTACAAGAACCTCACGGTTCGCGAGGCCTG
||||||||||||||||||||||||||||||||||||||||||||||||||||||||||||

GCTTAACACCCTGGTGACCGCCGAGGTCATCTTCTGGTTCTACATCGGCGAGTGCATCGG
||||||||||||||||||||||||||||||||||||||||||||||||||||||||||||

CAAGCGTCACATTGTAGGCTACAATGTCTAAGCTTACTATAGTCTCCGCTTGGCAGTCAC
||||||||||||||||||||||||||||||||||||||||||||||||||||||||||||

TGGAATGGGCAACGTAATCCCTAACAGATGTGTATATTTATATGTCTGCGAACATTTCGA
||||||||||||||||||||||||||||||||||||||||||||||||||||||||||||

CTCTGAATAAAGTGAAATAGTAATTTAAAATTCCGAAAATTTTCGAAAAATACATTGTTT
||||||||||||||||||||||||||||||||||||||||||||||||||||||||||||

TTTGAAAACCGTTAGAACGTTTGCGCGGGATTTGTGTAAGCTAAAGATGAGGTGATGTAA
||||||||||||||||||||||||||||||||||||||||||||||||||||||||||||

AACCAAGTTTGGAATTAAAAGTTGACGATATTTAATGATAAAAATTAAAAAAATATATGT
||||||||||||||||||||||||||||||||||||||||||||||||||||||||||||

AATGTTATACATCAAATGTTTATGAAACGGTGTCGTAATCAAAGAGGCTAATGGTTCAGA
||||||||||||||||||||||||||||||||||||||||||||||||||||||||||||

AATACATAATATACTTAGAGCATTTAAAAGCACTCAAGAATAATTTTATTTAAAAAAAAA
||||||||||||||||||||||||||||||||||||||||||||||||||||||||||||

                          Upstream of ao start codon
AATAAATAAATCTAAATTTGCTTTTCATAGATAATTCATTTACACATTTTTTTAAACAAA
||||||||||||||||||||||||||||||||||||||||||||||||||||||||||||

GTAAAGTAAGATGTATTGAATTATTTTATTTATAAATAACGTTTTTTATTTGAATCTTGA
||||||||||||||||||||||||||||||||||||||||||||||||||||||||||||

AAAGCATTAATATTATTATTATTATACTTATATTTTTTAACAACAAAACTTTTGTAGACA
||||||||||||||||||||||||||||||||||||||||||||||||||||||||||||

GAGTAAATTTTTGTAATCTAACTGCGGTCACACTTTACTTTAGTTACCTTTCGATCGGAA
|||||||||||||||||||||||||||||||||||||||||||||||||| |||||||||

                                          3XP3 promoter sequence
GAAGAACCCGGCTGGATCTAATTCAATTAGAGACTAATTCAATTAGAGCTAATTCAATTA
|||||||||||||||||

GGATCCAAGCTTATCGATTTCGAACCCTCGACCGCCGGAGTATAAATAGAGGCGCTTCGT

CTACGGAGCGACAATTCAATTCAAACAAGCAAAGTGAACACGTCGCTAAGCGAAAGCTAA

GCAAATAAACAAGCGCAGCTGAACAAGCTAAACAATCGGCTCGAAGCCGGTCGCCACCAT

                     dsRed
GGCCTCCTCCGAGGACGTCATCAAGGAGTTCATGCGCTTCAAGGTGCGCATGGAGGGCTC

CGTGAACGGCCACGAGTTCGAGATCGAGGGCGAGGGCGAGGGCCGCCCCTACGAGGGCAC

CCAGACCGCCAAGCTGAAGGTGACCAAGGGCGGCCCCCTGCCCTTCGCCTGGGACATCCT

GTCCCCCCAGTTCCAGTACGGCTCCAAGGTGTACGTGAAGCACCCCGCCGACATCCCCGA

CTACAAGAAGCTGTCCTTCCCCGAGGGCTTCAAGTGGGAGCGCGTGATGAACTTCGAGGA

CGGCGGCGTGGTGACCGTGACCCAGGACTCCTCCCTCCAGGACGGCTCCTTCATCTACAA

GGTGAAGTTCATCGGCGTGAACTTCCCCTCCGACGGCCCCGTAATGCAGAAGAAGACTAT

GGGCTGGGAGGCGTCCACCGAGCGCCTGTACCCCCGCGACGGCGTGCTGAAGGGCGAGAT

CCACAAGGCCCTGAAGCTGAAGGACGGCGGCCACTACCTGGTGGAGTTCAAGTCCATCTA

CATGGCCAAGAAGCCCGTGCAGCTGCCCGGCTACTACTACGTGGACTCCAAGCTGGACAT

CACCTCCCACAACGAGGACTACACCATCGTGGNNCAGTACGAGCGCGCCGAGGGCCGCCA

                         Downstream of ao Ter codon
CCACCTGTTCCTGTAGATCTCGTTTTACCTTTTCAGTAATGTCTTTATTTACAATGATAA
                |||||||| ||||||||||||||| ||||||||||||||| |||

GAGAACATTTCTTTGTTTTATTTAATCAAAGACTGTTAATATTCCAAGTACTGTTTAAAT
|  |||||||||||||||||||||||||||||||||||||||||||||||||||||||||

CTGACAAACACTTTTTAATTCGACTTTGCTATATTGTATTGAAGGCCCACTTCAAACTTG
||||||||||||||||||||||||||||||||||||||||||||||||||||||||||||

AGGCCCAATCCGACGTTTAAAGTTTCATGTAACATTGTATCTCGACTGCGACTACGTTGA
||||||||||||||||||||||||||||||||||||||||||||||||||||||||||||

AGATTGGTGGTGACCGTCCTCGTTGTATTCCTTGTTGACCTTAAGGGAATGATTAATAAC
||||||||||||||||||||||||||||||||||||||||||||||||||||||||||||

GCTCGAGCCGCCTGGAGGATAGTTCGGGGCAGGCCCTGANCCCGGGCCGCTGGCAGGCTG
||||||||||||||||||||||||||||||||||||||| ||||||||||||||||||||

CTGTGCCTGAGGCTGATTGGGGATCGCATTTGGGTTGTTAACTGGATTCTGTCCGGGAGC
|||||||||| |||||||||||||||||||||||||||||||||||||||||||||||||
```

**Figure S2**
